# Are pangolins the intermediate host of the 2019 novel coronavirus (2019-nCoV) ?

**DOI:** 10.1101/2020.02.18.954628

**Authors:** Ping Liu, Jing-Zhe Jiang, Xiu-Feng Wan, Yan Hua, Xiaohu Wang, Fanghui Hou, Jing Chen, Jiejian Zou, Jinping Chen

## Abstract

The outbreak of 2019-nCoV pneumonia (COVID-19) in the city of Wuhan, China has resulted in more than 70,000 laboratory confirmed cases, and recent studies showed that 2019-nCoV (SARS-CoV-2) could be of bat origin but involve other potential intermediate hosts. In this study, we assembled the genomes of coronaviruses identified in sick pangolins. The molecular and phylogenetic analyses showed that pangolin Coronaviruses (pangolin-CoV) are genetically related to both the 2019-nCoV and bat Coronaviruses but do not support the 2019-nCoV arose directly from the pangolin-CoV. Our study also suggested that pangolin be natural host of *Betacoronavirus*, with a potential to infect humans. Large surveillance of coronaviruses in pangolins could improve our understanding of the spectrum of coronaviruses in pangolins. Conservation of wildlife and limits of the exposures of humans to wildlife will be important to minimize the spillover risks of coronaviruses from wild animals to humans.

## Introduction

In December 2019, there was an outbreak of pneumonia with an unknown cause in Wuhan, Hubei province in China, with an epidemiological link to the Huanan Seafood Wholesale Market, which is a live animal and seafood market. Clinical presentations of this disease greatly resembled viral pneumonia. Through deep sequencing on the lower respiratory tract samples of patients, a novel coronavirus named the 2019 novel coronavirus (2019-nCoV) was identified [1]. Within less than 2 months, the viruses have spread to all provinces across China and 23 additional countries. As of February 19, 2020, the epidemic has resulted in 72,532 laboratory confirmed cases, 1,872 of which were fatal. With nearly three weeks of locking down Wuhan (and followed by many other cities across China), the toll of new cases and deaths are still rising.

To effectively control the diseases and prevent new spillovers, it is critical to identify the animal origin of this newly emerging coronavirus. In this wet market of Wuhan, high viral loads were reported in the environmental samples. However, variety of animals, including some wildlife, were sold on this market, and the number and species were very dynamics. It remains unclear which animal initiated the first infections.

Coronaviruses cause respiratory and gastrointestinal tract infections and are genetically classified into four major genera: *Alphacoronavirus*, *Betacoronavirus*, *Gammacoronavirus*, and *Deltacoronavirus*. The former two genera primarily infect mammals, whereas the latter two predominantly infect birds [2]. In addition to the 2019-nCoV, *Betacoronavirus* caused the 2003 SARS (severe acute respiratory syndrome) outbreaks and the 2012 MERS (Middle East respiratory syndrome) outbreaks in humans [3, 4]. Both SARS-CoV and MERS-CoV are of bat origin, but palm civets were shown to be an intermediate host for SARS-CoV [5] and dromedary camels for MERS-CoV [6].

Approximate 30-thousand-base genome of coronavirus codes up to 11 proteins, and the surface glycoprotein S protein binds to receptors on the host cell, initiating virus infection. Different coronaviruses can use distinct host receptors due to structural variations in the receptor binding domains of the virus S protein. SARS-CoV uses angiotensin-converting enzyme 2 (ACE2) as one of the main receptors [7] with CD209L as an alternative receptor [8], whereas MERS-CoV uses dipeptidyl peptidase 4 (DPP4, also known as CD26) as the primary receptor. Computational modeling analyses suggested that, similar to SARS-CoV, the 2019-nCoV uses ACE2 as the receptor [9].

Not soon after the release of the 2019-nCoV genome, a scientist released a full genome of a coronavirus, Bat-CoV-RaTG13, from bat (*Rhinolophus sinicus*), which is colonized in Yunan province, nearly 2,000 km away from Wuhan. Bat-CoV-RaTG13 was 96% identical at the whole genome level to the 2019-nCoV, suggesting the 2019-nCoV, could be of bat origin [10]. However, with rare direct contacts between such bats and humans, similar to SARS-CoV and MERS-CoV, it seems to be more likely that the spillover of 2019-nCoV to humans from another intermediate host rather than directly from bats.

The goal of this study is to determine the genetic relationship between a coronavirus from two groups of sick pangolins and the 2019-nCoV and to assess whether pangolins could be a potential intermediate host for the 2019-nCoV.

## Results

In March of 2019, we detected *Betacoronavirus* in three animals from two sets of smuggling *Malayan* pangolins *(Manis javanica*) (n=26) intercepts by Guangdong customs [11]. All three animals suffered from serious respiratory diseases and failed to be rescued by the Guangdong Wildlife Rescue Center [11] (Table S2). Through metagenomic sequencing and *de novo* assembling, we recovered 38 contigs ranging from 380 to 3,377 nucleotides, and the nucleotide sequence identities among the contigs from these three samples were 99.54%. Thus, we pooled sequences from three samples and assembled the draft genome of this pangolin origin coronavirus, so called pangolin-CoV-2020 (Accession No.: GWHABKW00000000), which was approximately 29,380 nucleotides, with approximately 84% coverage of the virus genome (Figure 1a).

**Figure 1.**
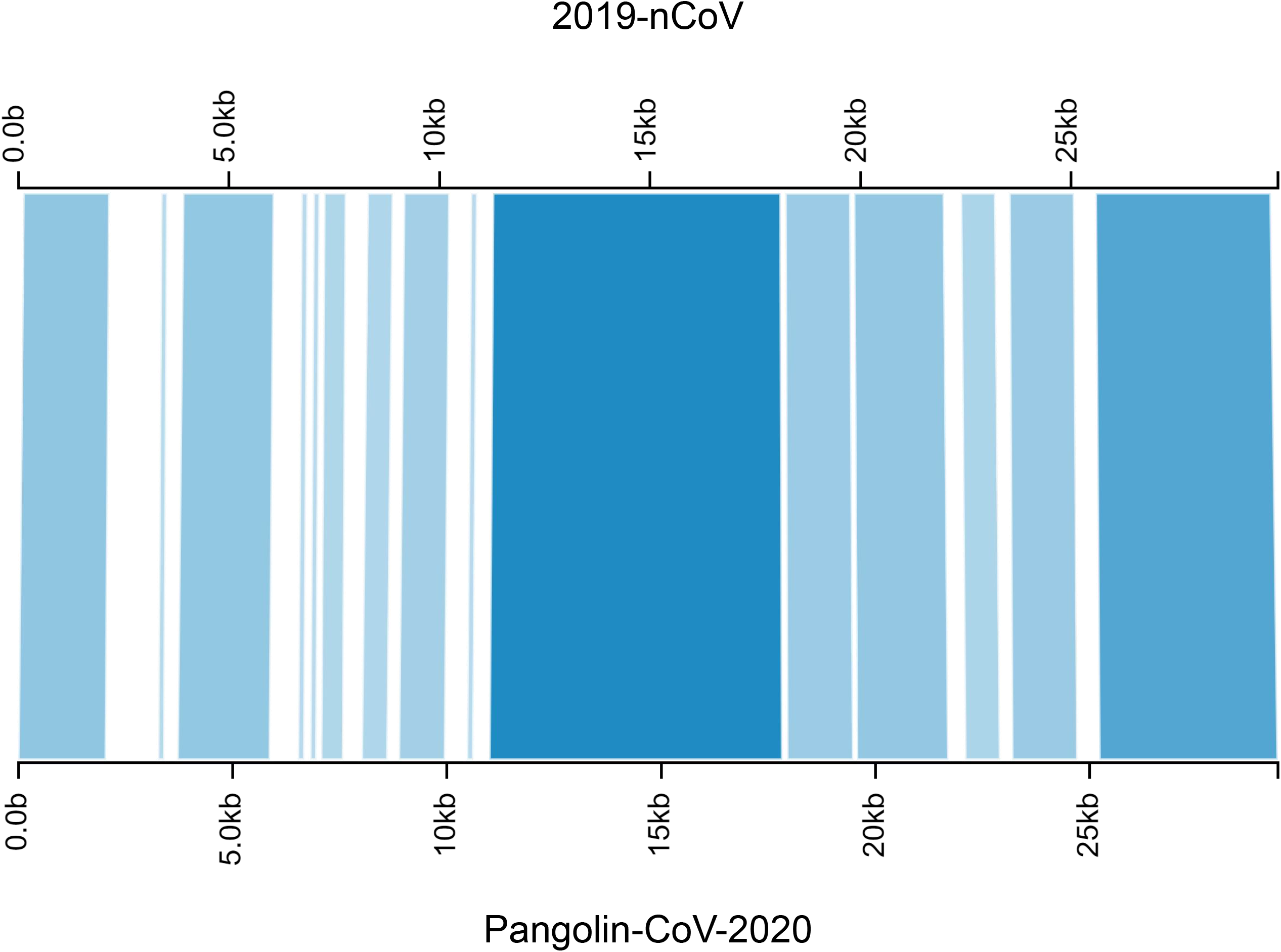
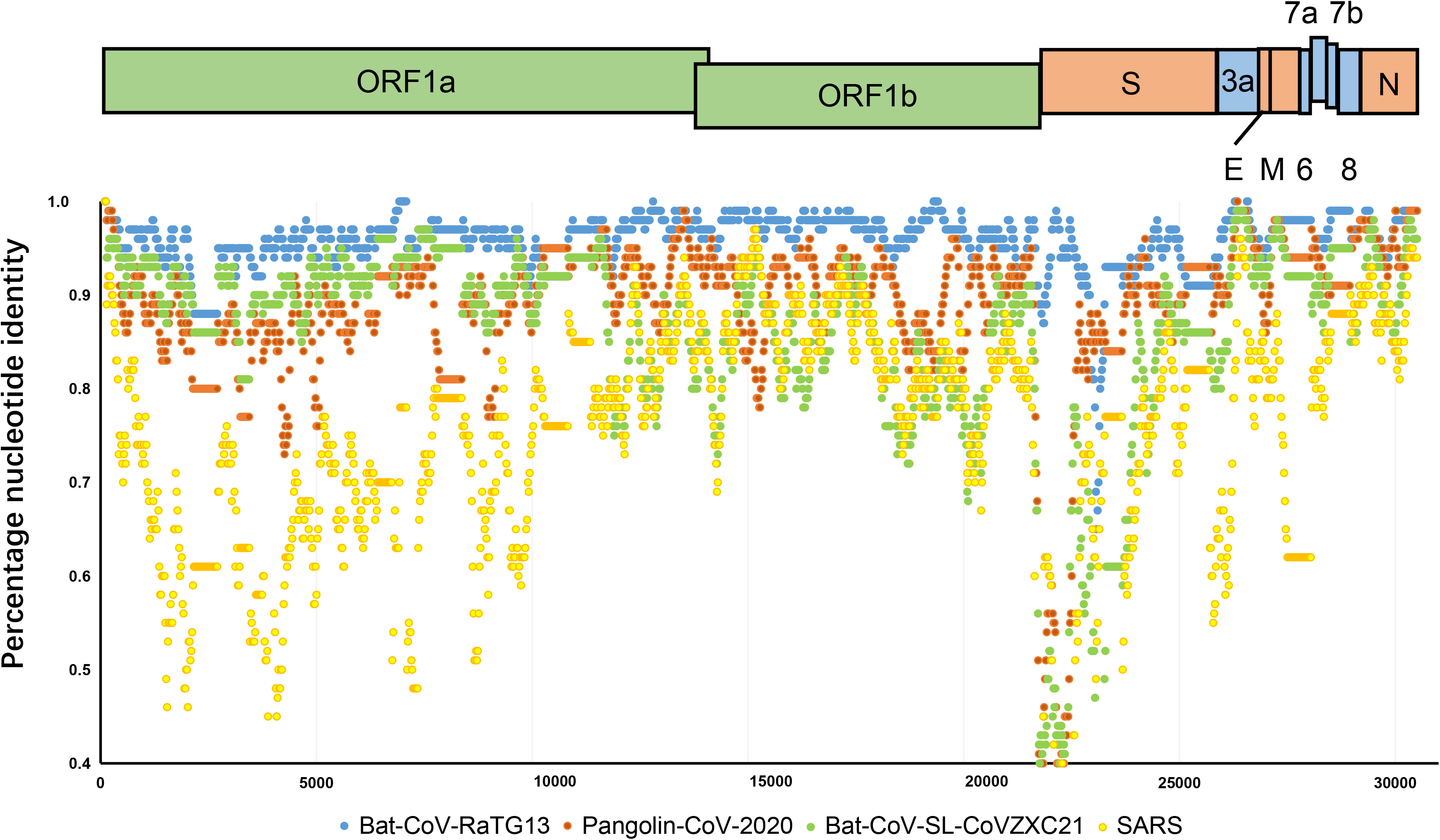
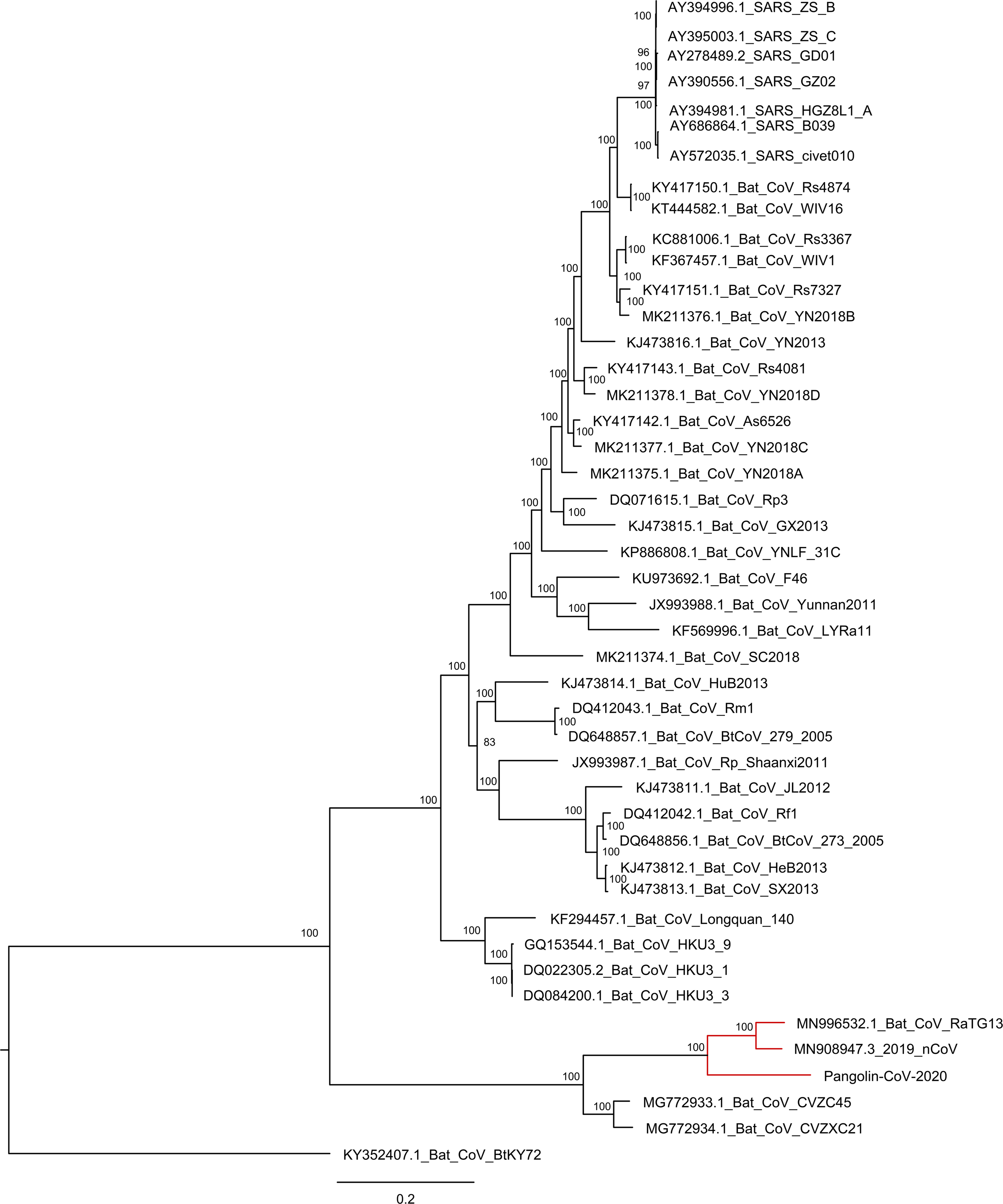
Genomic comparison of pangolin-CoV-2020, 2019-nCoV, and other coronaviruses. a) genomic alignment of pangolin-CoV-2020 and 2019-nCoV, white indicates missing sequence; b) Similarity plot based on the full-length genome sequence of 2019-nCoV. Full-length genome sequences of Bat-CoV-RaTg13, Bat-CoV-SL-CoVZXC21, SARS, and pangolin-CoV-2020 draft genome were used as subject sequences; c) Phylogenetic tree based on nucleotide sequences of complete genomes of coronaviruses.

Strikingly, genomic analyses suggested the pangolin-CoV-2020 has a high identity with both 2019-nCoV and Bat-CoV-RaTG13, the proposed origin of the 2019-nCoV [10] (Figure 1b; Figure 1c). The nucleotide sequence identity between pangolin-CoV-2020 and 2019-nCoV was 90.23% whereas the protein sequence identities for individual proteins can be up to 100% (Table S3; Table S4). The nucleotide sequence between pangolin-CoV-2020 and Bat-CoV-RaTG13 was 90.15% whereas that for the corresponding regions between 2019-nCoV and Bat-CoV-RaTG13 was 96.12% (Table 1).

**Table 1.**
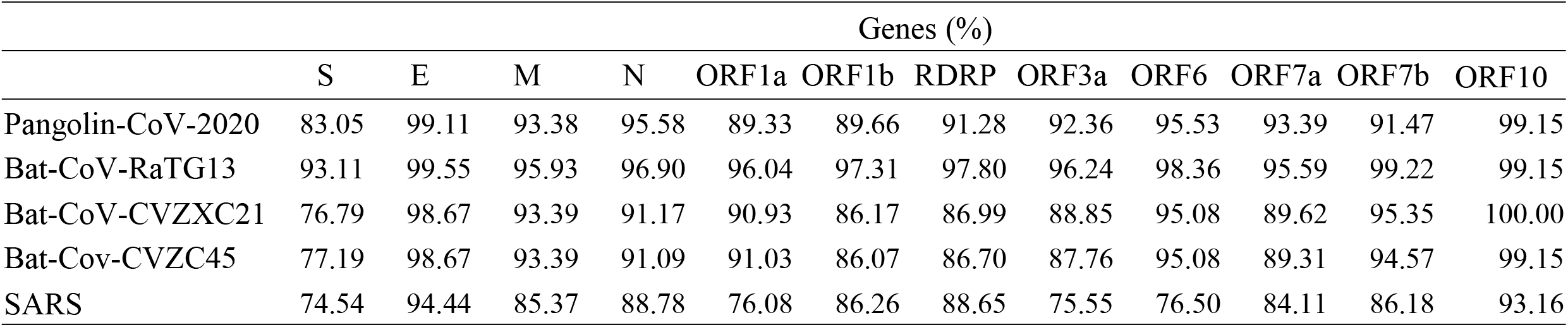
Nucleotide sequence identities among the genes of pangolin-CoV-2020 and other representative coronavirus against 2019 nCoV.

The nucleotide sequence identities of the surface glycoprotein Spike (S) protein genes between pangolin-CoV-2020 and 2019-nCoV was 82.21%, and the Bat-CoV-RaTG13 and 2019-nCoV shared the highest sequence identity of 92.59% (Table 1). There was a low similarity of 72.63% between the S genes of pangolin-CoV-2020 and SARS-CoV. Nucleotide sequence analyses suggested the S gene was relatively more genetic diverse in the S1 region than the S2 region (Figure 2a). Furthermore, the S proteins of pangolin-CoV-2020 and 2019-nCoV had a sequence identity of 89.78% (Table 2), sharing a very conserved receptor binding motif (RBM) (Figure S1), which is more conserved than in Bat-CoV-RaTG13. These results support that pangolin-CoV-2020 and 2019-nCoV, and SARS-CoV could all share the same receptor ACE2. The presence of highly identical RBMs in pangolin-CoV-2020 and 2019-nCoV means that this motif was likely already present in the virus before jumping to humans. However, it is interesting that both pangolin-CoV-2020 and Bat-CoV-RaTG13 lack a S1/S2 cleavage site (~680-690 aa) whereas 2019-nCoV possess (Figure S1).

**Table 2.**
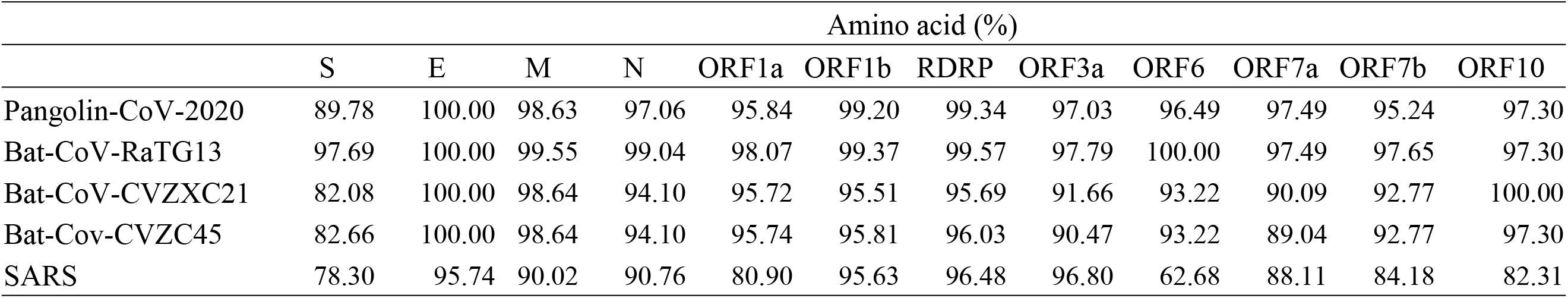
Protein sequence identities among the genes of pangolin-CoV-2020 and other representative coronavirus against 2019 nCoV.

**Figure 2.**
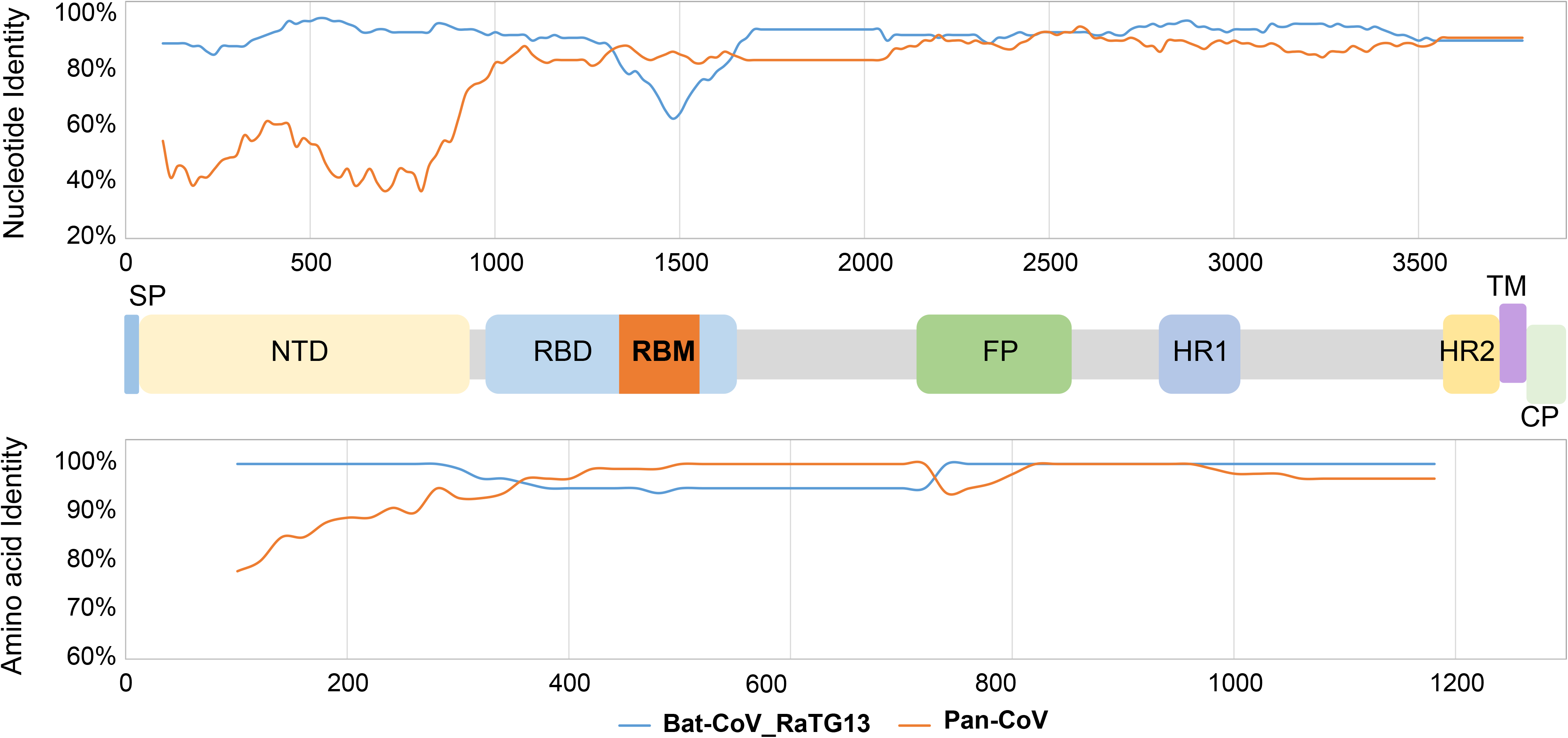
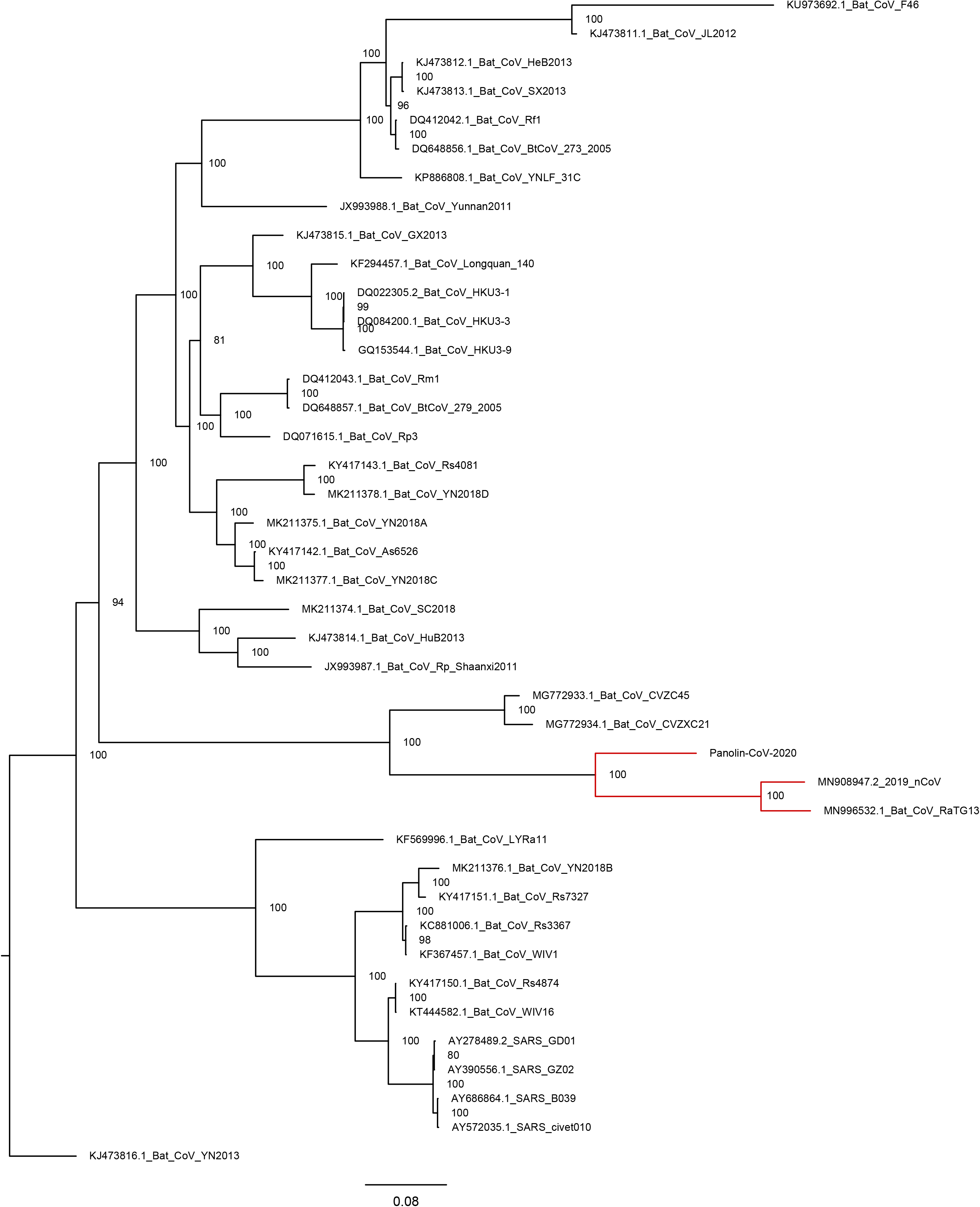
Genetic analyses of the spike surface glycoprotein of pangolin-CoV-2020, 2019-nCoV, and other coronaviruses. a) similarity plot based on the spike surface glycoprotein amino acid and nucleotide sequence of 2019-nCoV. Bat-CoV-RaTg13, and pangolin-CoV-2020 were used as subject sequences; b) phylogenetic tree of S genes.

Phylogenetic analyses suggested that the S genes of pangolin-CoV-2020, 2019-nCoV and three bat origin coronaviruses (Bat-CoV-RaTG13, Bat-CoV-CVZXC21, and Bat-Cov-CVZC45) were genetically more similar to each other than other viruses in the same family (Figure 2b). The S gene of Bat-CoV-RaTG13 was genetically closer to each other than pangolin-CoV-2020, Bat-CoV-CVZXC21, and Bat-Cov-CVZC45. Similar tree topologies were observed for the RdRp gene and other genes (Figure 3a-d; Figure S2a-h). At the whole genomic level, the 2019-nCoV is also genetically closer to Bat-CoV-RaTG13 than pangolin-CoV-2020 (Figure 1c).

**Figure 3.**
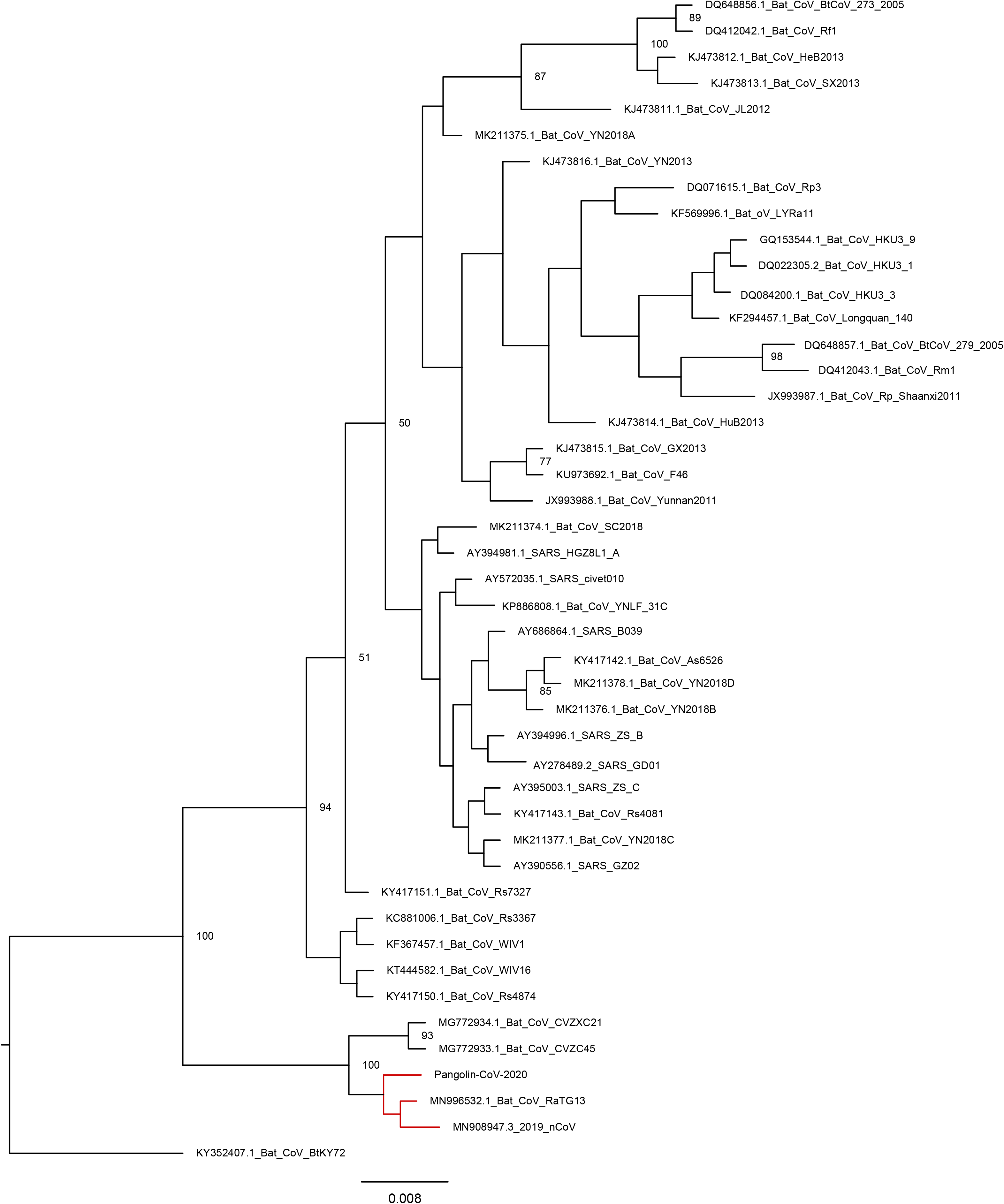
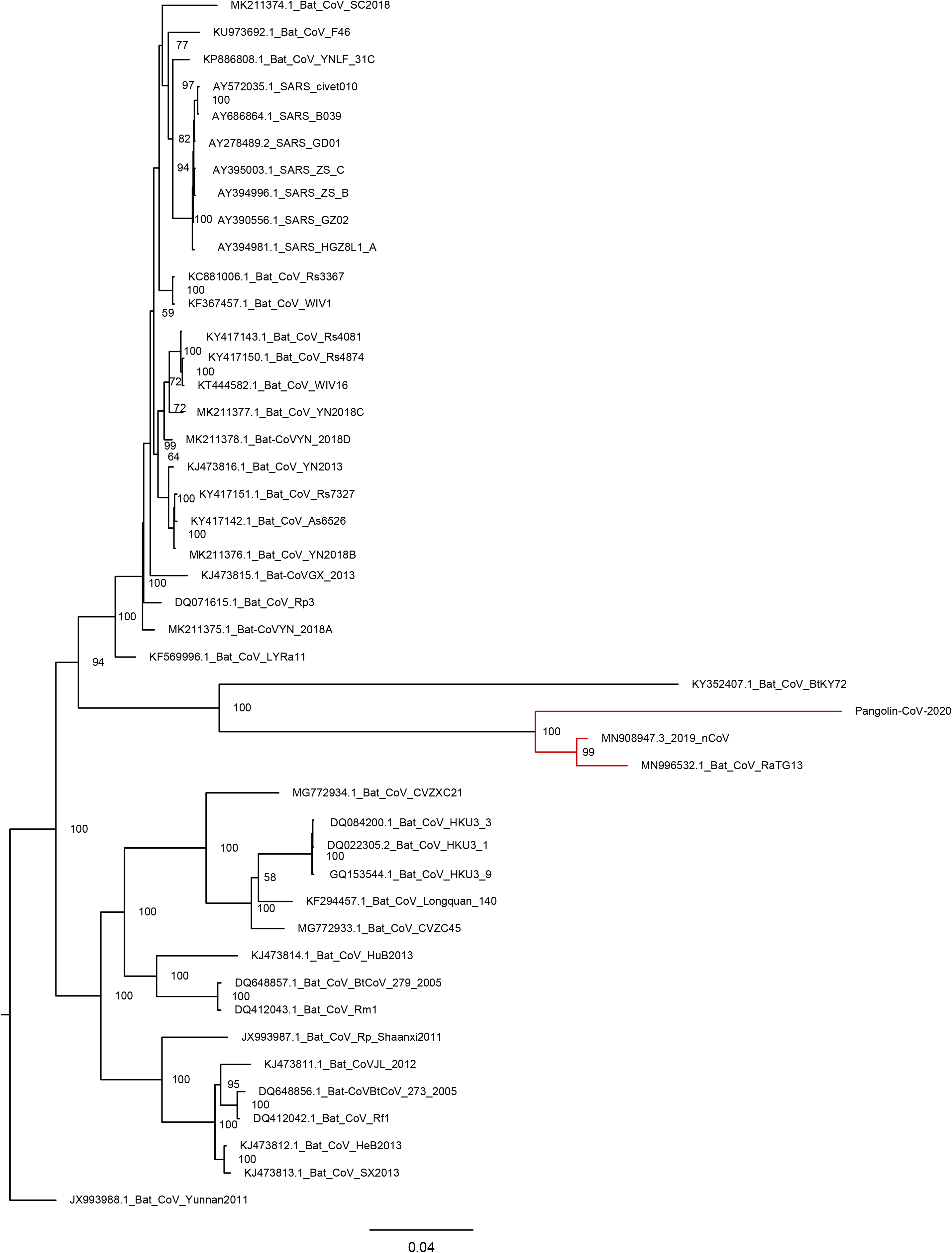
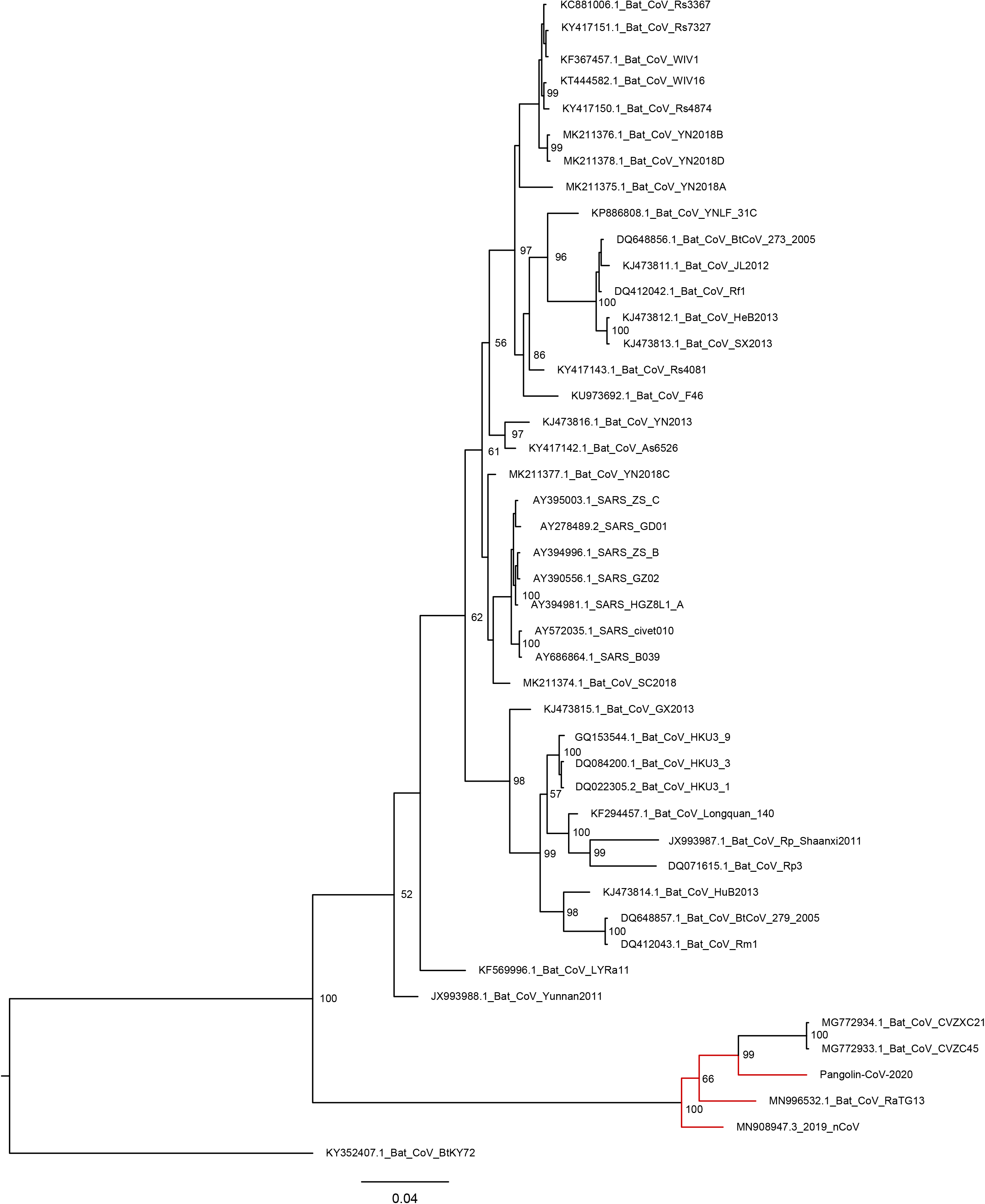
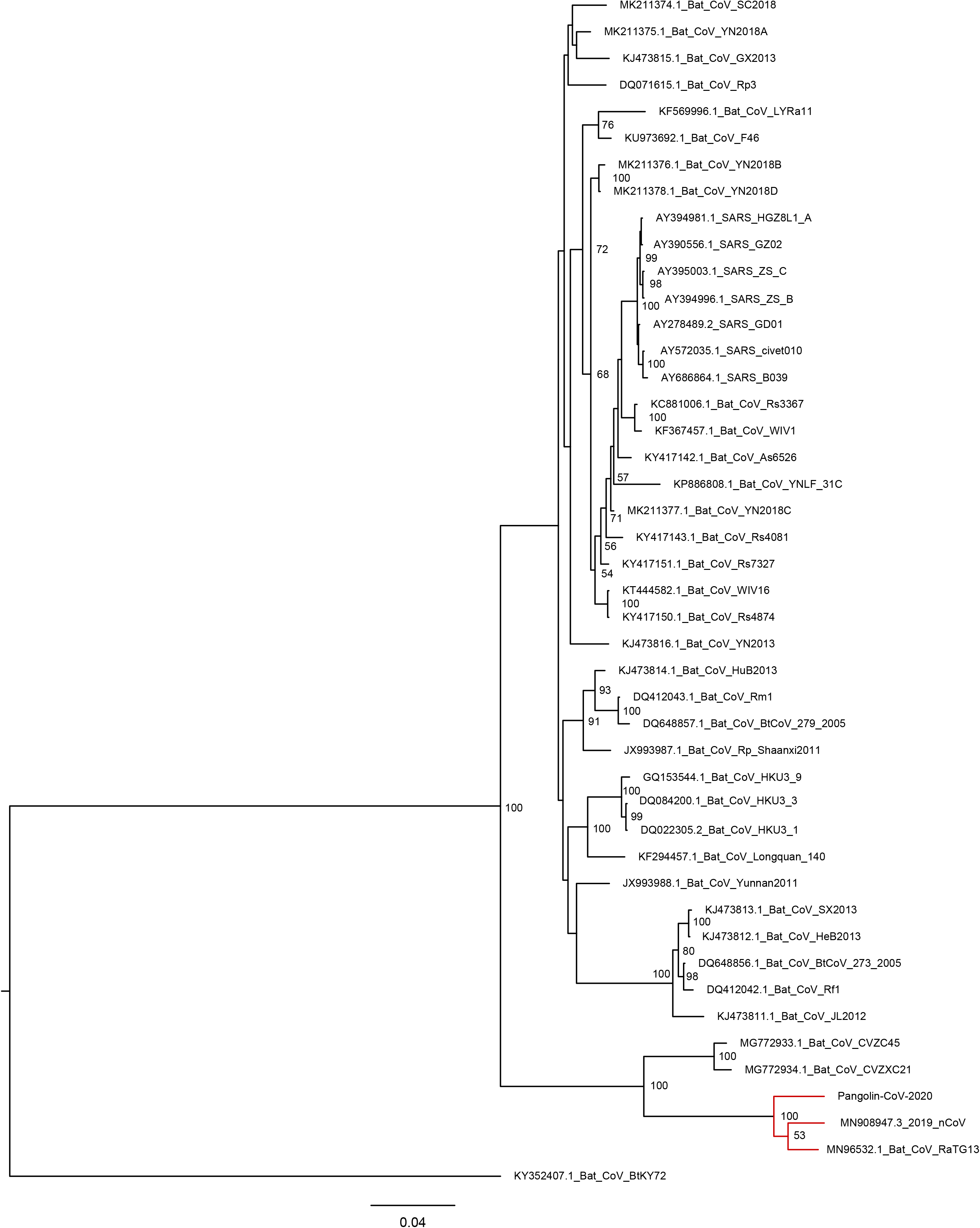
Phylogenetic analyses of a) small envelope gene, b) RNA-dependent RNA polymerase (RdRp) gene, c) matrix protein, and d) nucleocapsid protein sequences of coronaviruses from different hosts.

## Discussion

In this study, we assembled the genomes of coronaviruses identified in sick pangolins and our results showed that a pangolin coronavirus (pangolin-CoV-2020) is genetically associated with both 2019-nCoV and a group of bat coronaviruses. There is a high sequence similarity between pangolin-CoV-2020 and 2019-nCoV. However, phylogenetic analyses did not support the 2019-nCoV arose directly from the pangolin-CoV-2020.

It is of interest that the genomic sequences for coronaviruses detected from two batches of pangolins intercepted by two different customs at different dates were all be associated with bat coronaviruses. The reads from the third pangolin acquired in July of 2019 were relatively less abundant than the two from the first two pangolin samples acquired in March of 2019. Although we are unclear whether these two batches of smuggling and exotic pangolins were from the same origin, our results indicated that pangolin be a natural host for *Betacoronavirus*es, which could be enzootic in pangolins. All three exotic pangolins detected with *Betacoronavirus*es in this study were very sick with serious respiratory diseases and failed to be rescued. However, these pangolins were very stressful in the transportation freight when being intercepted by the customs. It is unclear whether this coronavirusis a common virus flora in the respiratory tracts of pangolins. Nevertheless, the pathogenesis of this coronavirus to pangolin remains to be studied.

Compared to the genomic sequence of pangolin-CoV-2020 we assembled in this study, phylogenetic trees suggested that a bat origin coronavirus (i.e., Bat-CoV-RaTG13) was more genetically close to the 2019-nCoV at both individual gene and genomic sequence level. Interestingly, the cleavage site between S1 and S2 at the 2019-nCoV had multiple insertions (i.e. PRRA), compared to that of Bat-CoV-RaTG13 and pangolin-CoV-2020, which were similar. Thus, although it is clear that 2019-nCoV is of bat origin, it is likely another intermediate host could be involved in emergence of the 2019-nCoV.

The S protein of coronaviruses bind to host receptors via receptor-binding domains (RBDs), and plays an essential role in initiating virus infection and determines host tropism [2]. A prior study suggested that the 2019-nCoV, SARS-CoV, and Bat-CoV-RaTG13 had similar RBDs, suggested all of them use the same receptor ACE2 [9]. Our analyses showed that pangolin-CoV-2020 had a very conserved RBD to these three viruses rather than MERS-CoV, suggesting that pangolin-CoV is very likely to use ACE2 as its receptor. On the other hand, ACE2 receptor is present in pangolins with a high sequence conservation with those in the gene homolog in humans. However, the zoonosis of this pangolin-CoV-2020 remains unclear.

The host range of animal origin coronaviruses was promiscuous [12]. It is critical to determine the natural reservoir and the host range of coronaviruses, especially their potential of causing zoonosis. In the last two decades, besides the 2019-nCoV, SARS and MERS caused serious outbreaks in humans, lead to thousands of deaths [3, 4, 13, 14]. Although all of three zoonotic coronaviruses were shown to be of bat origin, they seemed to use different intermediate hosts. For example, farmed palm civets were suggested to be an intermediate host for SARS to be spilled over to humans although the details on how to link bat and farmed palm civets are unclear [15, 16, 17]. Most recently, dromedary camels in Saudi Arabia were shown to harbor three different coronavirus species, including a dominant MERS-CoV lineage that was responsible for the outbreaks in the Middle East and South Korea during 2015 [18]. Although this present study does not support pangolins would be an intermediate host for the emergence of the 2019-nCoV, our results do not prevent the possibility that other CoVs could be circulating in pangolins. Thus, large surveillance of coronaviruses in the pangolins could improve our understanding the spectrum of coronaviruses in the pangolins. Conservation of wildlife and limits of the exposures of humans to wildlife will be important to minimize the spillover risks coronaviruses from wild animals to humans.

In summary, this study suggested pangolins be a natural host of *Betacoronaviru*s, with an unknown potential to infect humans. However, our data do not support the 2019-nCoV evolved directly from the pangolin-CoV.

## Materials and Methods

### Data selection

During our routine wildlife rescue efforts, one of the goals was to identify pathogens causing wildlife diseases. In 2019, we were involved in two events of pangolin rescues, one involved with 21 smuggling pangolins in March and the second with 6 smuggling pangolins in July. From those pangolins failed to be rescued, we collected samples from different tissues and subjected for metagenomic analyses. Through viral metagenomics analyses of lower respiratory tract samples from these pangolins, we detected coronavirus in three individual animals [11]. Two of these animals were from the first batch of Malayan pangolins intercepted by Meizhou, Yangjiang, and Jiangmen customs, and the third one was from the second batch in a freight being transported from Qingyuan to Heyuan. The RNA samples from these three individuals were subjected to deep sequencing. To determine the read abundance of coronaviruses in each sample, we mapped clean reads without ribosomes and host sequences to an in-house virus reference data separated from the GenBank non-redundant nucleotide database.

### Genomic assembly and sequence analyses

After examining the high similarity among the samples from three animals, to maximize the coverage of the virus genome, clean reads from three animals were pooled together and *de novo* assembled using MEGAHIT v1.2.9 [19]. The assembled contigs were used as references for mapping those the rest unmapped reads using Salmon v0.14.1 [20], and multiple rounds were implemented to maximize the mapping (Table S2).

A total of 38 contigs were identified to be highly similar to the 2019-nCoV genome (accession MN908947.3) using BLASTn and tBLASTx. GapFiller v1.10 and SSPACE v3.0 were used to fill gaps and draft pangolin-CoV-2020 genome was constructed with ABACAS v1.3.1 (http://abacas.sourceforge.net/) [21, 22, 23].

Multiple sequence alignments were conducted using CLUSTAL Ov1.2.4 [24]. Simplot analyses were conducted with SimPlot v3.5.1 to determine the sequence similarity among 2019-nCoV (MN908947.3), pangolin-CoV-2020, Bat-CoV-RaTG13 (MN996532.1), and SARS-CoV (AY395003.1) at both the genomic sequence level and at individual gene level [25]. Sequence identity was calculated utilizing p-diatance in MEGA v10.1.7 [26].

### Phylogenetic analyses

We downloaded 44 full-length genome sequences of coronaviruses isolated from different hosts from the public database (Table S1). Phylogenetic analyses were performed based on their whole genome sequences, encoding ORFs of RNA-dependent RNA polymerase (RdRp gene), the receptor binding protein spike protein (S gene), small envelope protein (E gene), as well as all other gene sequences were conducted utilizing Mrbayes [27] with 50,000,000 generations and the 25% of the generations as burnin. Best models were determined by jModeltest v2.1.7 [28]. Then, all the trees were visualized and exported as vector diagrams with FigTree v1.4.4 (http://tree.bio.ed.ac.uk/software/figtree/).

## Acknowledgements

We thank the De-Chun Lin and Tao Jin from Magigene Biotech. and Hanghui Kong from South China Botanical Garden support for bioinformatics analysis. This project was supported by wildlife disease monitoring and early warning system maintenance project from National Forestry and Grassland Administration (2019072), GDAS Special Project of Science and Technology Development (grant number 2020GDASYL-20200103090, 2018GDASCX-0107),Guangzhou Science Technology and Innovation Commission (grant number 201804020080), Natural Science Foundation of China (grant number 31972847), Guangzhou science and technology project (grant number 2019001), and 2019-nCoV wildlife origin project from Guangdong department of science and technology.

## Supplementary Information

### Supplementary Tables

**Table S1.** Accession numbers and strain IDs of coronaviruses strains isolated from different hosts.

**Table S2.** Number of sequencing reads assigned to different viruses in each pangolin sample. We only focus on Coronaviruses in this study.

**Table S3.** The blast results for the assembled nucleotide contigs of pangolin-CoV-2020 and 2019-nCoV.

**Table S4.** The blast results for the translated proteins from assembled nucleotide contigs of pangolin-CoV-2020 and 2019-nCoV.

### Supplementary Figures

**Figure S1.** Amino acid sequence alignment of the spike surface glycoprotein of the Pangolin-CoV-2020 with 2019-nCoV and Bat-CoV-RaTG13.

**Figure S2. Phylogenetic analyses of** a) ORF1a, b) ORF1b, c) ORF3a, d) ORF6, e) ORF7a, f) ORF7b, g) ORF8, and h) ORF10 gene sequences of coronaviruses from different hosts.

