## supplemental files for "Are pangolins the intermediate host of the 2019 novel coronavirus (2019-nCoV) ?"

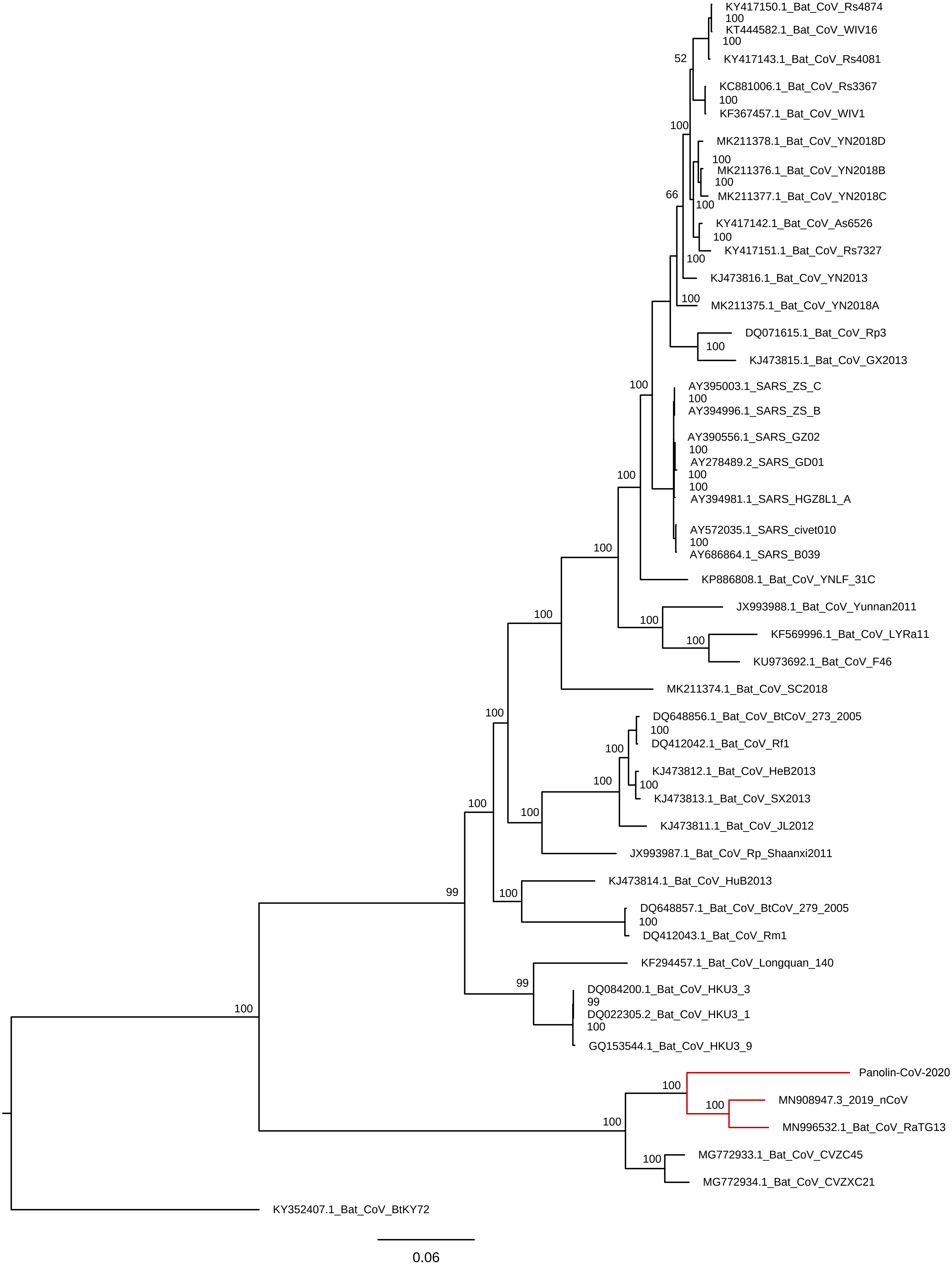
KY417150.1_Bat_CoV_Rs4874
KT444582.1_Bat_CoV_WIV16
KY417143.1_Bat_CoV_Rs4081
KC881006.1_Bat_CoV_Rs3367
KF367457.1_Bat_CoV_WIV1
MK211378.1_Bat_CoV_YN2018D
MK211376.1_Bat_CoV_YN2018B
MK211377.1_Bat_CoV_YN2018C
KY417142.1_Bat_CoV_As6526
KY417151.1_Bat_CoV_Rs7327
KJ473816.1_Bat_CoV_YN2013
MK211375.1_Bat_CoV_YN2018A
DQ071615.1_Bat_CoV_Rp3
KJ473815.1_Bat_CoV_GX2013
AY395003.1_SARS_ZS_C
AY394996.1_SARS_ZS_B
AY390556.1_SARS_GZ02
AY278489.2_SARS_GD01
AY394981.1_SARS_HGZ8L1_A
AY572035.1_SARS_civet010
AY686864.1_SARS_B039
KP886808.1_Bat_CoV_YNLF_31C
JX993988.1_Bat_CoV_Yunnan2011
KF569996.1_Bat_CoV_LYRa11
KU973692.1_Bat_CoV_F46
MK211374.1_Bat_CoV_SC2018
DQ648856.1_Bat_CoV_BtCoV_273_2005
DQ412042.1_Bat_CoV_Rf1
KJ473812.1_Bat_CoV_HeB2013
KJ473813.1_Bat_CoV_SX2013
KJ473811.1_Bat_CoV_JL2012
JX993987.1_Bat_CoV_Rp_Shaanxi2011
KJ473814.1_Bat_CoV_HuB2013
DQ648857.1_Bat_CoV_BtCoV_279_2005
DQ412043.1_Bat_CoV_Rm1
KF294457.1_Bat_CoV_Longquan_140
DQ084200.1_Bat_CoV_HKU3_3
DQ022305.2_Bat_CoV_HKU3_1
GQ153544.1_Bat_CoV_HKU3_9
Panolin-CoV-2020
MN908947.3_2019_nCoV
MN996532.1_Bat_CoV_RaTG13
MG772933.1_Bat_CoV_CVZC45
MG772934.1_Bat_CoV_CVZXC21
KY352407.1_Bat_CoV_BtKY72
0.06

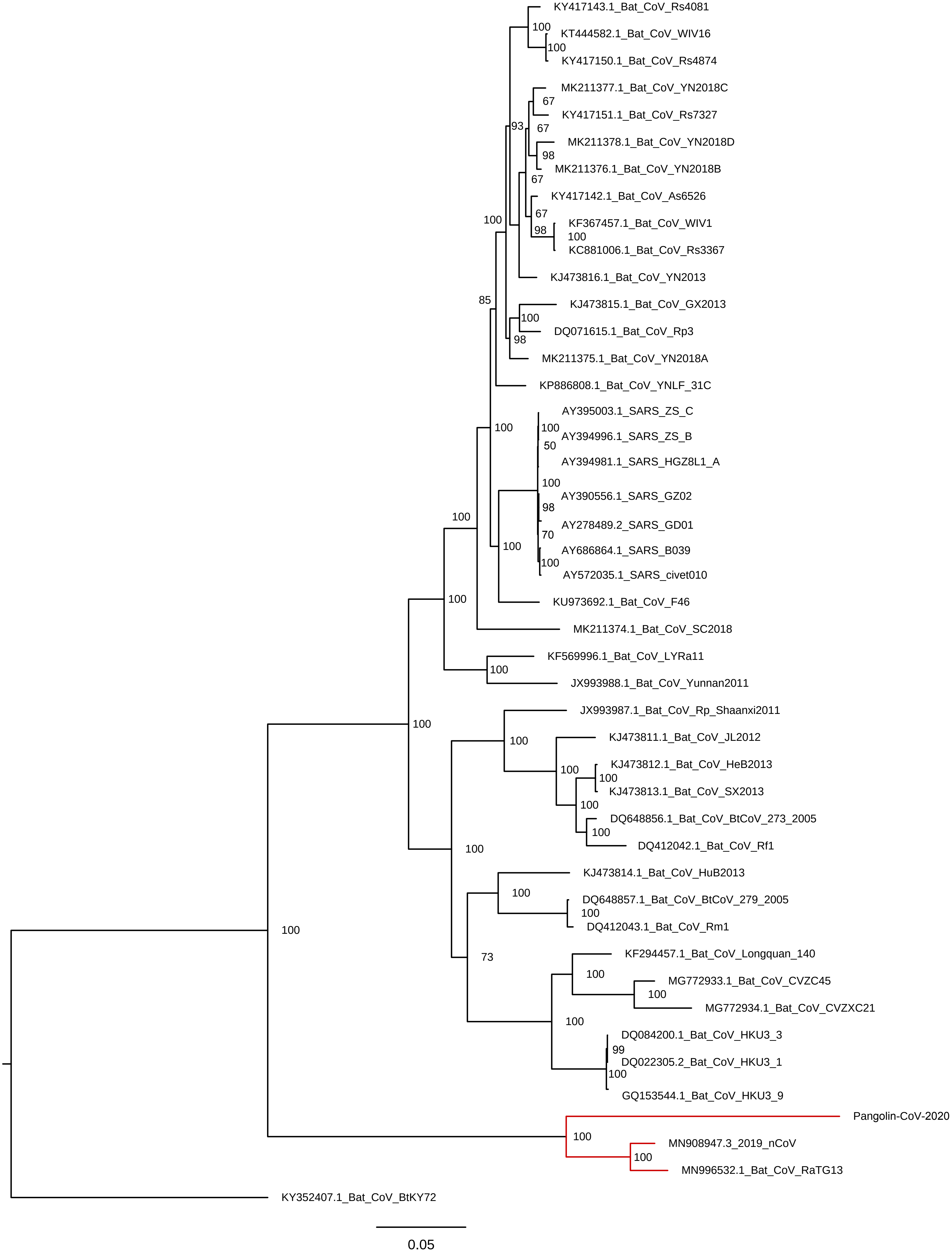
KY417143.1_Bat_CoV_Rs4081
KT444582.1_Bat_CoV_WIV16
KY417150.1_Bat_CoV_Rs4874
MK211377.1_Bat_CoV_YN2018C
KY417151.1_Bat_CoV_Rs7327
MK211378.1_Bat_CoV_YN2018D
MK211376.1_Bat_CoV_YN2018B
KY417142.1_Bat_CoV_As6526
KF367457.1_Bat_CoV_WIV1
KC881006.1_Bat_CoV_Rs3367
KJ473816.1_Bat_CoV_YN2013
KJ473815.1_Bat_CoV_GX2013
DQ071615.1_Bat_CoV_Rp3
MK211375.1_Bat_CoV_YN2018A
KP886808.1_Bat_CoV_YNLF_31C
AY395003.1_SARS_ZS_C
AY394996.1_SARS_ZS_B
AY394981.1_SARS_HGZ8L1_A
AY390556.1_SARS_GZ02
AY278489.2_SARS_GD01
AY686864.1_SARS_B039
AY572035.1_SARS_civet010
KU973692.1_Bat_CoV_F46
MK211374.1_Bat_CoV_SC2018
KF569996.1_Bat_CoV_LYRa11
JX993988.1_Bat_CoV_Yunnan2011
JX993987.1_Bat_CoV_Rp_Shaanxi2011
KJ473811.1_Bat_CoV_JL2012
KJ473812.1_Bat_CoV_HeB2013
KJ473813.1_Bat_CoV_SX2013
DQ648856.1_Bat_CoV_BtCoV_273_2005
DQ412042.1_Bat_CoV_Rf1
KJ473814.1_Bat_CoV_HuB2013
DQ648857.1_Bat_CoV_BtCoV_279_2005
DQ412043.1_Bat_CoV_Rm1
KF294457.1_Bat_CoV_Longquan_140
MG772933.1_Bat_CoV_CVZC45
MG772934.1_Bat_CoV_CVZXC21
DQ084200.1_Bat_CoV_HKU3_3
DQ022305.2_Bat_CoV_HKU3_1
GQ153544.1_Bat_CoV_HKU3_9
Pangolin-CoV-2020
MN908947.3_2019_nCoV
MN996532.1_Bat_CoV_RaTG13
KY352407.1_Bat_CoV_BtKY72
0.05

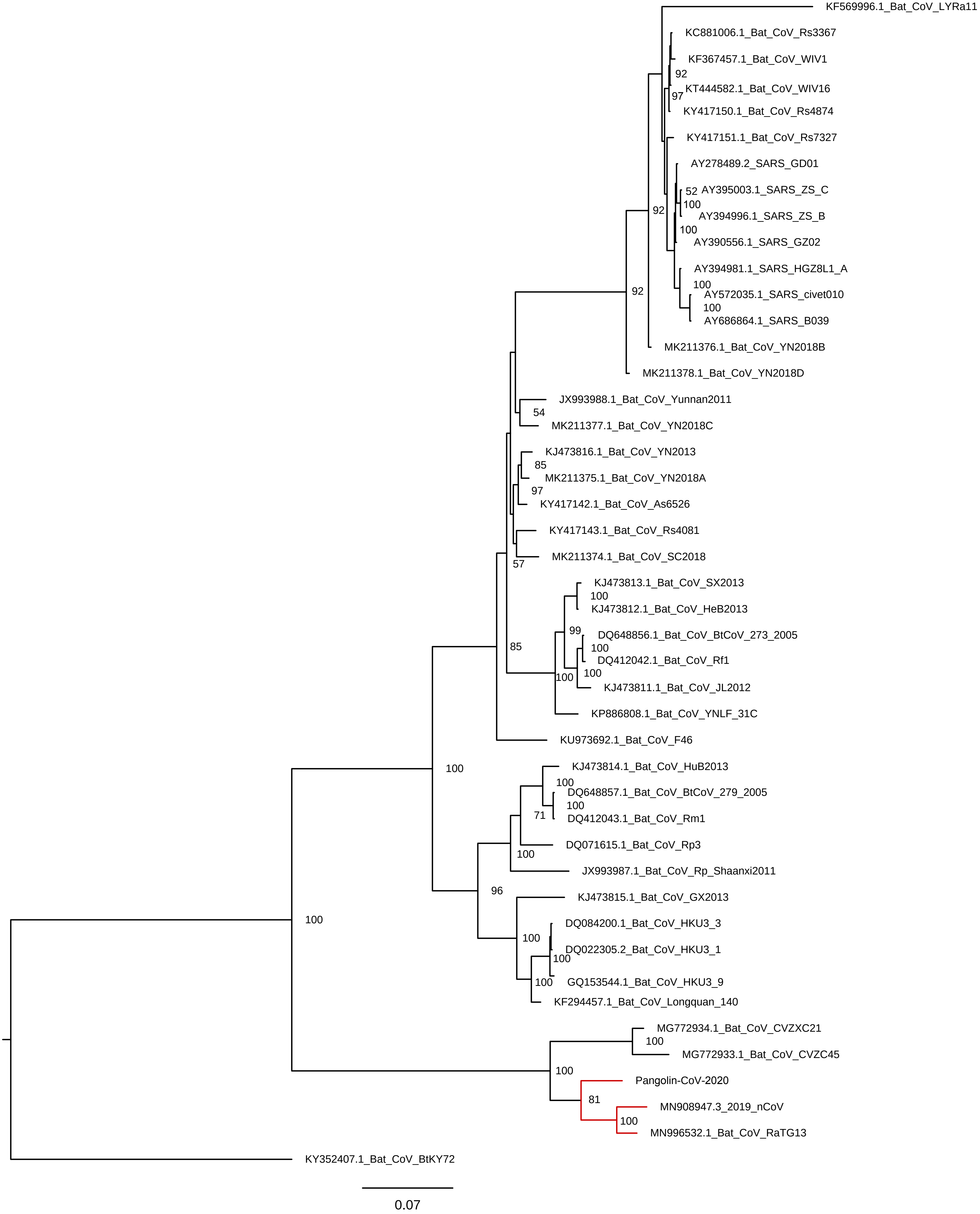
KF569996.1_Bat_CoV_LYRa11
KC881006.1_Bat_CoV_Rs3367
KF367457.1_Bat_CoV_WIV1
92
KT444582.1_Bat_CoV_WIV16
97
KY417150.1_Bat_CoV_Rs4874
KY417151.1_Bat_CoV_Rs7327
AY278489.2_SARS_GD01
52
AY395003.1_SARS_ZS_C
100
92
AY394996.1_SARS_ZS_B
100
AY390556.1_SARS_GZ02
AY394981.1_SARS_HGZ8L1_A
100
92
AY572035.1_SARS_civet010
100
AY686864.1_SARS_B039
MK211376.1_Bat_CoV_YN2018B
MK211378.1_Bat_CoV_YN2018D
JX993988.1_Bat_CoV_Yunnan2011
54
MK211377.1_Bat_CoV_YN2018C
KJ473816.1_Bat_CoV_YN2013
85
MK211375.1_Bat_CoV_YN2018A
97
KY417142.1_Bat_CoV_As6526
KY417143.1_Bat_CoV_Rs4081
MK211374.1_Bat_CoV_SC2018
57
KJ473813.1_Bat_CoV_SX2013
100
KJ473812.1_Bat_CoV_HeB2013
99
DQ648856.1_Bat_CoV_BtCoV_273_2005
85
100
DQ412042.1_Bat_CoV_Rf1
100
100
KJ473811.1_Bat_CoV_JL2012
KP886808.1_Bat_CoV_YNLF_31C
KU973692.1_Bat_CoV_F46
KJ473814.1_Bat_CoV_HuB2013
100
100
DQ648857.1_Bat_CoV_BtCoV_279_2005
100
71
DQ412043.1_Bat_CoV_Rm1
DQ071615.1_Bat_CoV_Rp3
100
JX993987.1_Bat_CoV_Rp_Shaanxi2011
96
KJ473815.1_Bat_CoV_GX2013
100
DQ084200.1_Bat_CoV_HKU3_3
100
DQ022305.2_Bat_CoV_HKU3_1
100
100
GQ153544.1_Bat_CoV_HKU3_9
KF294457.1_Bat_CoV_Longquan_140
MG772934.1_Bat_CoV_CVZXC21
100
MG772933.1_Bat_CoV_CVZC45
100
Pangolin-CoV-2020
81
MN908947.3_2019_nCoV
100
MN996532.1_Bat_CoV_RaTG13
KY352407.1_Bat_CoV_BtKY72
0.07

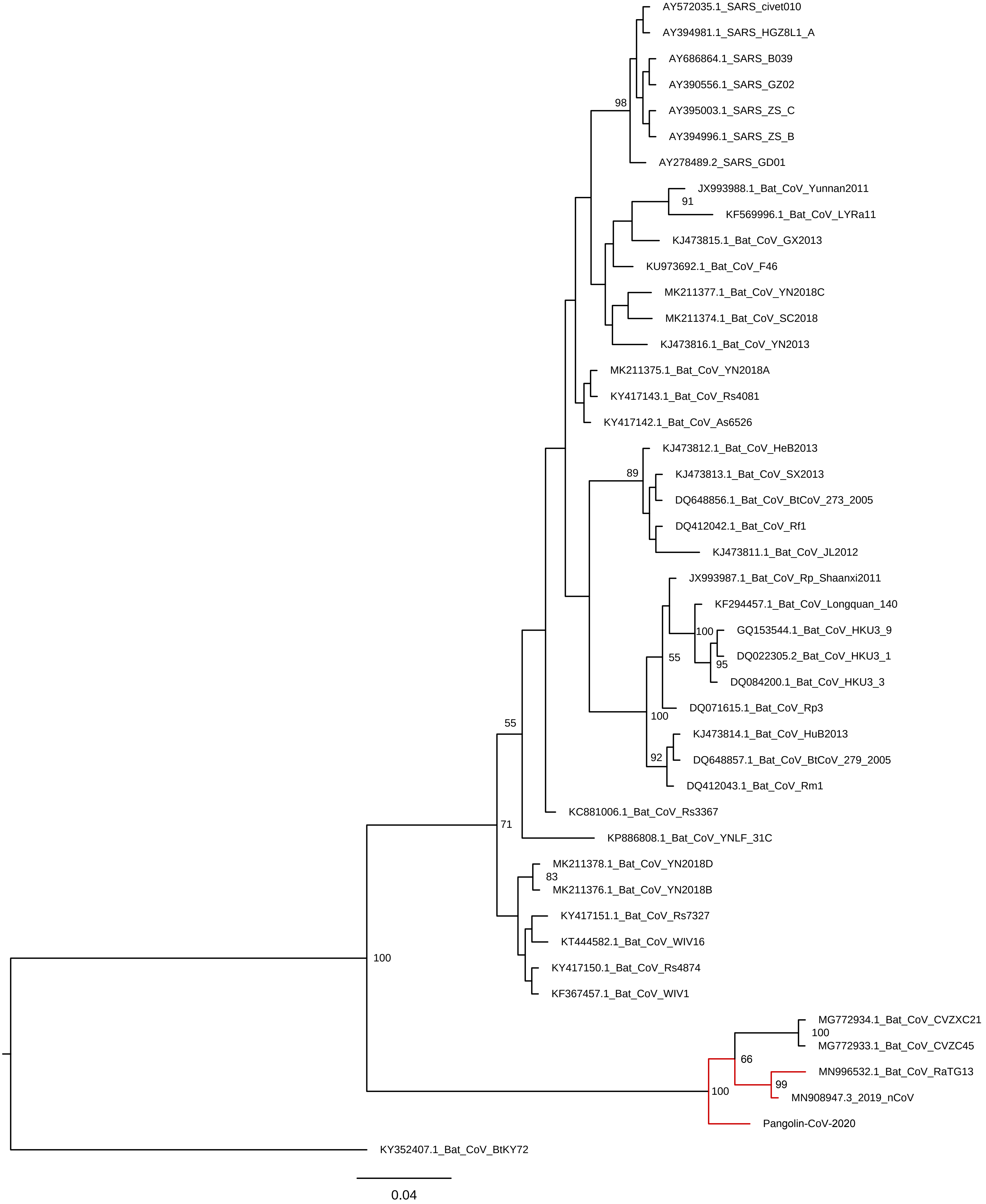
AY572035.1_SARS_civet010
AY394981.1_SARS_HGZ8L1_A
AY686864.1_SARS_B039
AY390556.1_SARS_GZ02
AY395003.1_SARS_ZS_C
AY394996.1_SARS_ZS_B
AY278489.2_SARS_GD01
98
JX993988.1_Bat_CoV_Yunnan2011
91
KF569996.1_Bat_CoV_LYRa11
KJ473815.1_Bat_CoV_GX2013
KU973692.1_Bat_CoV_F46
MK211377.1_Bat_CoV_YN2018C
MK211374.1_Bat_CoV_SC2018
KJ473816.1_Bat_CoV_YN2013
MK211375.1_Bat_CoV_YN2018A
KY417143.1_Bat_CoV_Rs4081
KY417142.1_Bat_CoV_As6526
KJ473812.1_Bat_CoV_HeB2013
89
KJ473813.1_Bat_CoV_SX2013
DQ648856.1_Bat_CoV_BtCoV_273_2005
DQ412042.1_Bat_CoV_Rf1
KJ473811.1_Bat_CoV_JL2012
JX993987.1_Bat_CoV_Rp_Shaanxi2011
KF294457.1_Bat_CoV_Longquan_140
100
GQ153544.1_Bat_CoV_HKU3_9
DQ022305.2_Bat_CoV_HKU3_1
95
DQ084200.1_Bat_CoV_HKU3_3
55
DQ071615.1_Bat_CoV_Rp3
100
KJ473814.1_Bat_CoV_HuB2013
92
DQ648857.1_Bat_CoV_BtCoV_279_2005
DQ412043.1_Bat_CoV_Rm1
55
KC881006.1_Bat_CoV_Rs3367
71
KP886808.1_Bat_CoV_YNLF_31C
MK211378.1_Bat_CoV_YN2018D
83
MK211376.1_Bat_CoV_YN2018B
KY417151.1_Bat_CoV_Rs7327
KT444582.1_Bat_CoV_WIV16
100
KY417150.1_Bat_CoV_Rs4874
KF367457.1_Bat_CoV_WIV1
MG772934.1_Bat_CoV_CVZXC21
100
MG772933.1_Bat_CoV_CVZC45
66
MN996532.1_Bat_CoV_RaTG13
99
100
MN908947.3_2019_nCoV
Pangolin-CoV-2020
KY352407.1_Bat_CoV_BtKY72
0.04

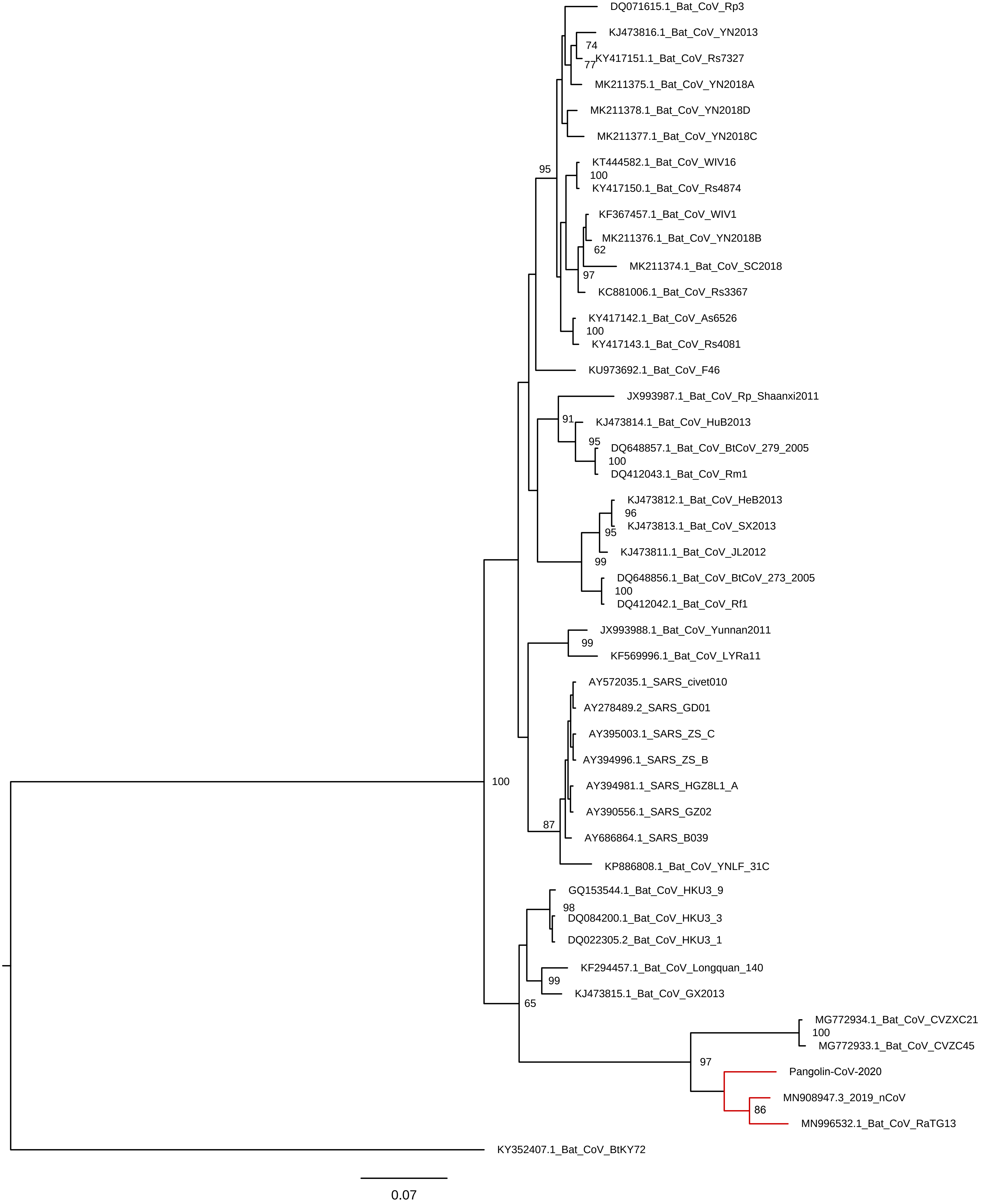

DQ071615.1_Bat_CoV_Rp3
KJ473816.1_Bat_CoV_YN2013
74
KY417151.1_Bat_CoV_Rs7327
77
MK211375.1_Bat_CoV_YN2018A
MK211378.1_Bat_CoV_YN2018D
MK211377.1_Bat_CoV_YN2018C
95
KT444582.1_Bat_CoV_WIV16
100
KY417150.1_Bat_CoV_Rs4874
KF367457.1_Bat_CoV_WIV1
MK211376.1_Bat_CoV_YN2018B
62
MK211374.1_Bat_CoV_SC2018
97
KC881006.1_Bat_CoV_Rs3367
KY417142.1_Bat_CoV_As6526
100
KY417143.1_Bat_CoV_Rs4081
KU973692.1_Bat_CoV_F46
JX993987.1_Bat_CoV_Rp_Shaanxi2011
91
KJ473814.1_Bat_CoV_HuB2013
95
DQ648857.1_Bat_CoV_BtCoV_279_2005
100
DQ412043.1_Bat_CoV_Rm1
KJ473812.1_Bat_CoV_HeB2013
96
KJ473813.1_Bat_CoV_SX2013
95
KJ473811.1_Bat_CoV_JL2012
99
DQ648856.1_Bat_CoV_BtCoV_273_2005
100
DQ412042.1_Bat_CoV_Rf1
JX993988.1_Bat_CoV_Yunnan2011
99
KF569996.1_Bat_CoV_LYRa11
AY572035.1_SARS_civet010
AY278489.2_SARS_GD01
AY395003.1_SARS_ZS_C
AY394996.1_SARS_ZS_B
100
AY394981.1_SARS_HGZ8L1_A
AY390556.1_SARS_GZ02
87
AY686864.1_SARS_B039
KP886808.1_Bat_CoV_YNLF_31C
GQ153544.1_Bat_CoV_HKU3_9
98
DQ084200.1_Bat_CoV_HKU3_3
DQ022305.2_Bat_CoV_HKU3_1
KF294457.1_Bat_CoV_Longquan_140
99
KJ473815.1_Bat_CoV_GX2013
65
MG772934.1_Bat_CoV_CVZXC21
100
MG772933.1_Bat_CoV_CVZC45
97
Pangolin-CoV-2020
MN908947.3_2019_nCoV
86
MN996532.1_Bat_CoV_RaTG13
KY352407.1_Bat_CoV_BtKY72
0.07

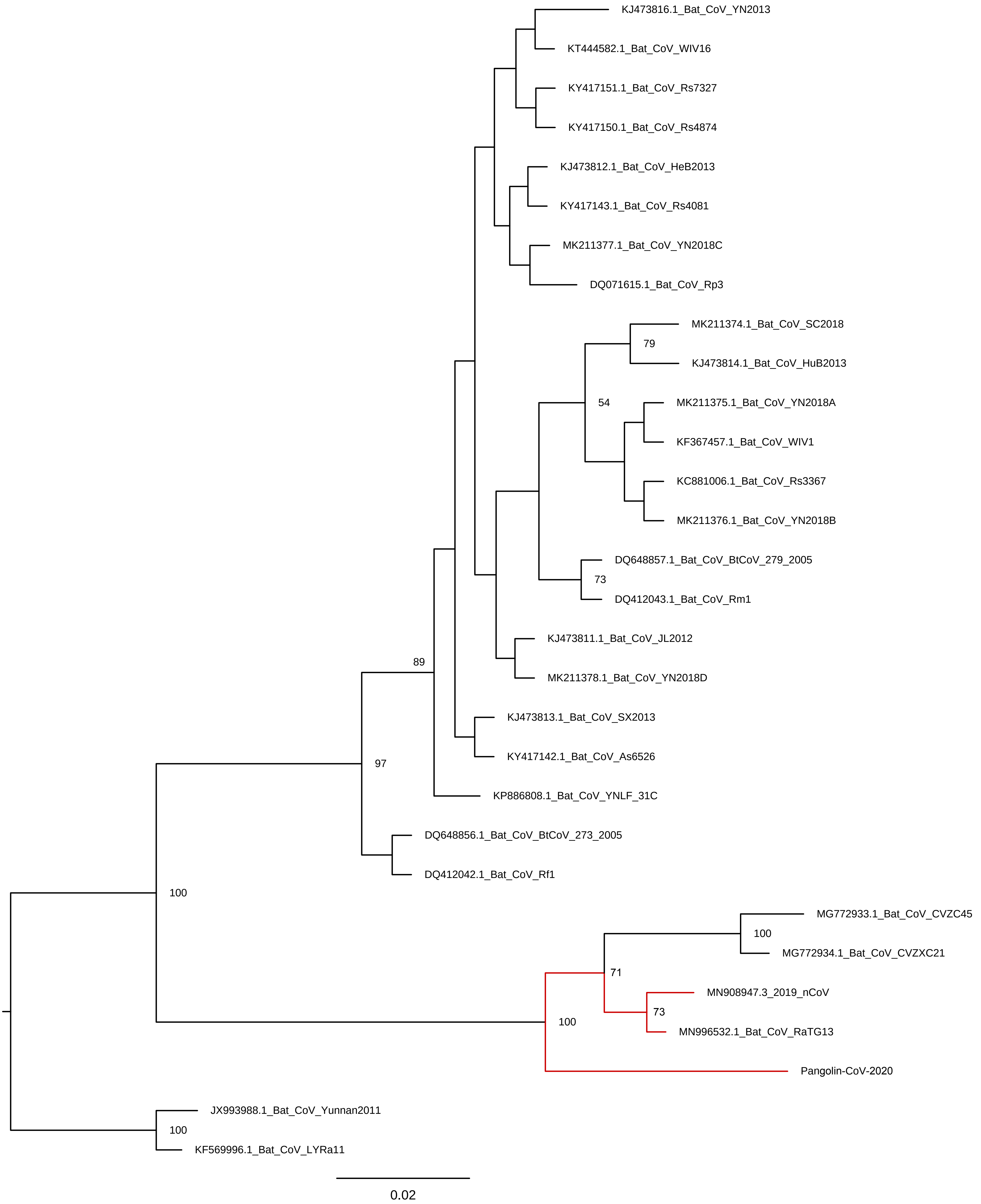
KJ473816.1_Bat_CoV_YN2013
KT444582.1_Bat_CoV_WIV16
KY417151.1_Bat_CoV_Rs7327
KY417150.1_Bat_CoV_Rs4874
KJ473812.1_Bat_CoV_HeB2013
KY417143.1_Bat_CoV_Rs4081
MK211377.1_Bat_CoV_YN2018C
DQ071615.1_Bat_CoV_Rp3
MK211374.1_Bat_CoV_SC2018
79
KJ473814.1_Bat_CoV_HuB2013
54
MK211375.1_Bat_CoV_YN2018A
KF367457.1_Bat_CoV_WIV1
KC881006.1_Bat_CoV_Rs3367
MK211376.1_Bat_CoV_YN2018B
DQ648857.1_Bat_CoV_BtCoV_279_2005
73
DQ412043.1_Bat_CoV_Rm1
KJ473811.1_Bat_CoV_JL2012
89
MK211378.1_Bat_CoV_YN2018D
KJ473813.1_Bat_CoV_SX2013
97
KY417142.1_Bat_CoV_As6526
KP886808.1_Bat_CoV_YNLF_31C
DQ648856.1_Bat_CoV_BtCoV_273_2005
DQ412042.1_Bat_CoV_Rf1
100
MG772933.1_Bat_CoV_CVZC45
100
MG772934.1_Bat_CoV_CVZXC21
71
MN908947.3_2019_nCoV
73
100
MN996532.1_Bat_CoV_RaTG13
Pangolin-CoV-2020
JX993988.1_Bat_CoV_Yunnan2011
100
KF569996.1_Bat_CoV_LYRa11
0.02

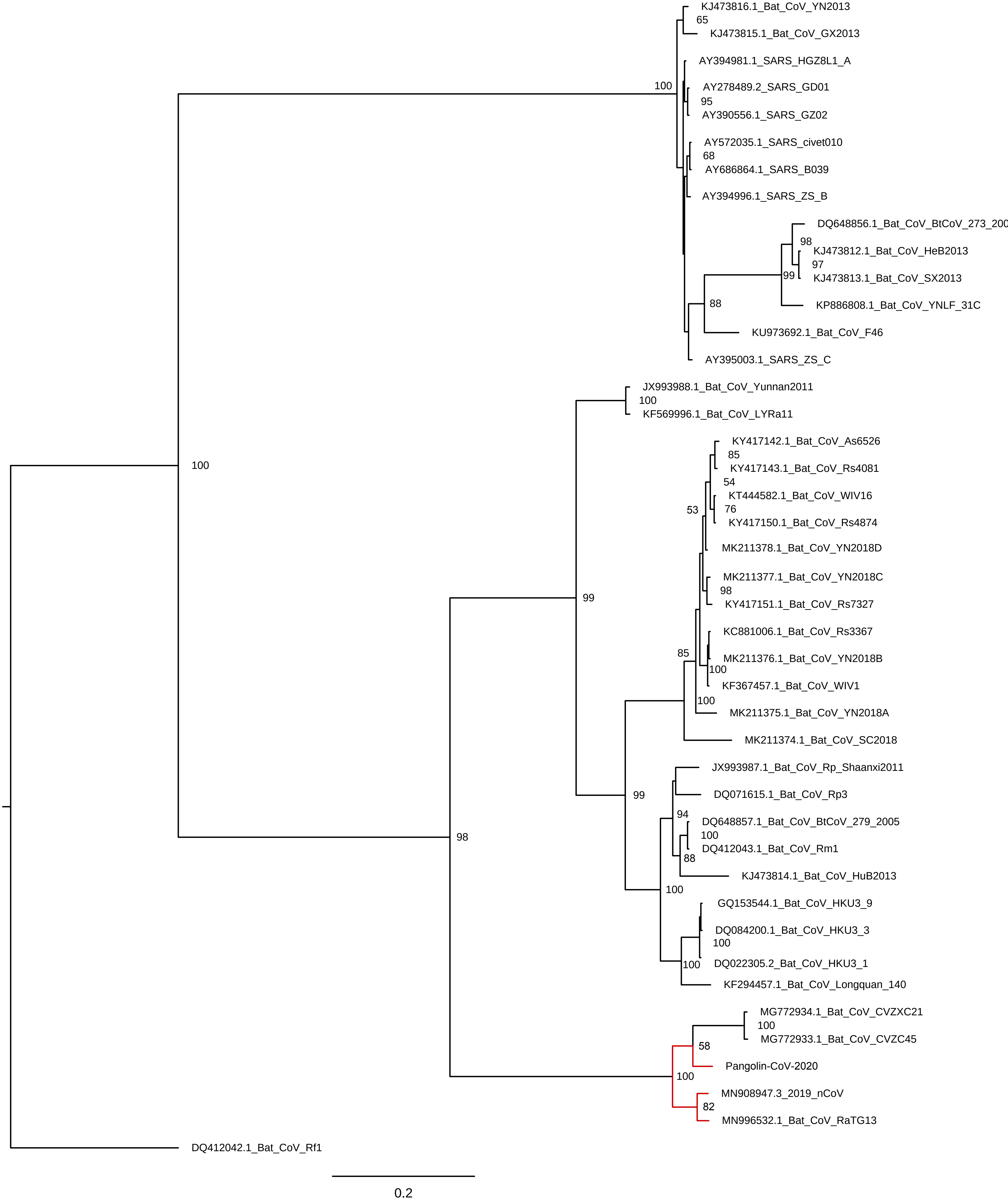
KJ473816.1_Bat_CoV_YN2013
65
KJ473815.1_Bat_CoV_GX2013
AY394981.1_SARS_HGZ8L1_A
100
AY278489.2_SARS_GD01
95
AY390556.1_SARS_GZ02
AY572035.1_SARS_civet010
68
AY686864.1_SARS_B039
AY394996.1_SARS_ZS_B
DQ648856.1_Bat_CoV_BtCoV_273_200
98
KJ473812.1_Bat_CoV_HeB2013
97
99
KJ473813.1_Bat_CoV_SX2013
88
KP886808.1_Bat_CoV_YNLF_31C
KU973692.1_Bat_CoV_F46
AY395003.1_SARS_ZS_C
JX993988.1_Bat_CoV_Yunnan2011
100
KF569996.1_Bat_CoV_LYRa11
KY417142.1_Bat_CoV_As6526
85
KY417143.1_Bat_CoV_Rs4081
54
KT444582.1_Bat_CoV_WIV16
53
76
KY417150.1_Bat_CoV_Rs4874
MK211378.1_Bat_CoV_YN2018D
MK211377.1_Bat_CoV_YN2018C
98
KY417151.1_Bat_CoV_Rs7327
KC881006.1_Bat_CoV_Rs3367
85
MK211376.1_Bat_CoV_YN2018B
100
KF367457.1_Bat_CoV_WIV1
100
MK211375.1_Bat_CoV_YN2018A
MK211374.1_Bat_CoV_SC2018
100
99
JX993987.1_Bat_CoV_Rp_Shaanxi2011
DQ071615.1_Bat_CoV_Rp3
99
94
DQ648857.1_Bat_CoV_BtCoV_279_2005
100
DQ412043.1_Bat_CoV_Rm1
88
98
KJ473814.1_Bat_CoV_HuB2013
100
GQ153544.1_Bat_CoV_HKU3_9
DQ084200.1_Bat_CoV_HKU3_3
100
100
DQ022305.2_Bat_CoV_HKU3_1
KF294457.1_Bat_CoV_Longquan_140
MG772934.1_Bat_CoV_CVZXC21
100
MG772933.1_Bat_CoV_CVZC45
58
Pangolin-CoV-2020
100
MN908947.3_2019_nCoV
82
MN996532.1_Bat_CoV_RaTG13
DQ412042.1_Bat_CoV_Rf1
0.2

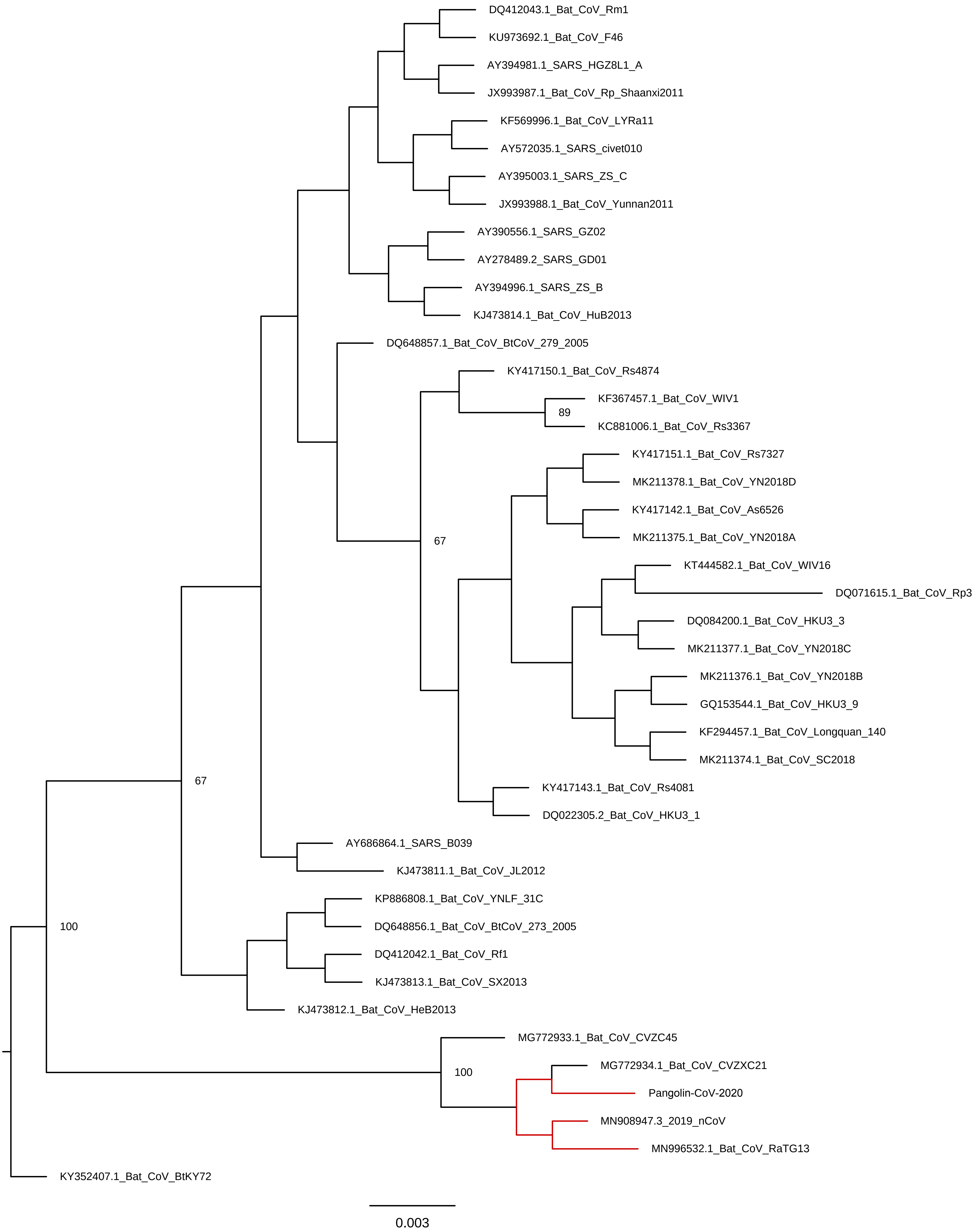
DQ412043.1_Bat_CoV_Rm1
KU973692.1_Bat_CoV_F46
AY394981.1_SARS_HGZ8L1_A
JX993987.1_Bat_CoV_Rp_Shaanxi2011
KF569996.1_Bat_CoV_LYRa11
AY572035.1_SARS_civet010
AY395003.1_SARS_ZS_C
JX993988.1_Bat_CoV_Yunnan2011
AY390556.1_SARS_GZ02
AY278489.2_SARS_GD01
AY394996.1_SARS_ZS_B
KJ473814.1_Bat_CoV_HuB2013
DQ648857.1_Bat_CoV_BtCoV_279_2005
KY417150.1_Bat_CoV_Rs4874
KF367457.1_Bat_CoV_WIV1
89
KC881006.1_Bat_CoV_Rs3367
KY417151.1_Bat_CoV_Rs7327
MK211378.1_Bat_CoV_YN2018D
KY417142.1_Bat_CoV_As6526
MK211375.1_Bat_CoV_YN2018A
67
KT444582.1_Bat_CoV_WIV16
DQ071615.1_Bat_CoV_Rp3
DQ084200.1_Bat_CoV_HKU3_3
MK211377.1_Bat_CoV_YN2018C
MK211376.1_Bat_CoV_YN2018B
GQ153544.1_Bat_CoV_HKU3_9
KF294457.1_Bat_CoV_Longquan_140
MK211374.1_Bat_CoV_SC2018
67
KY417143.1_Bat_CoV_Rs4081
DQ022305.2_Bat_CoV_HKU3_1
AY686864.1_SARS_B039
KJ473811.1_Bat_CoV_JL2012
KP886808.1_Bat_CoV_YNLF_31C
100
DQ648856.1_Bat_CoV_BtCoV_273_2005
DQ412042.1_Bat_CoV_Rf1
KJ473813.1_Bat_CoV_SX2013
KJ473812.1_Bat_CoV_HeB2013
MG772933.1_Bat_CoV_CVZC45
MG772934.1_Bat_CoV_CVZXC21
100
Pangolin-CoV-2020
MN908947.3_2019_nCoV
MN996532.1_Bat_CoV_RaTG13
KY352407.1_Bat_CoV_BtKY72
0.003

| Accession number | Strain ID |
|---|---|
| MN908947.3 | 2019-nCoV |
| MN996532.1 | Bat-CoV_RaTG13 |
| KJ473811.1 | Bat-CoV_JL2012 |
| MK211378.1 | Bat-CoV_YN2018D |
| KJ473815.1 | Bat-CoV_GX2013 |
| MK211375.1 | Bat-CoV_YN2018A |
| DQ648856.1 | Bat-CoV_BtCoV/273/2005 |
| DQ412042.1 | Bat-CoV_Rf1 |
| AY390556.1 | BARS_GZ02 |
| AY278489.2 | BARS_GD01 |
| AY572035.1 | BARS_civet010 |
| AY686864.1 | BARS_B039 |
| AY395003.1 | BARS_ZS-C |
| AY394996.1 | BARS_ZS-B |
| MK211376.1 | Bat-CoV_YN2018B |
| KC881006.1 | Bat-CoV_Rs3367 |
| KY417143.1 | Bat-CoV_Rs4081 |
| KU973692.1 | Bat-CoV_F46 |
| KJ473814.1 | Bat-CoV_HuB2013 |
| KY417151.1 | Bat-CoV_Rs7327 |
| KF367457.1 | Bat-CoV_WIV1 |
| KJ473816.1 | Bat-CoV_YN2013 |
| KP886808.1 | Bat-CoV_YNLF_31C |
| KJ473812.1 | Bat-CoV_HeB2013 |
| MK211377.1 | Bat-CoV_YN2018C |
| JX993987.1 | Bat-CoV_Rp/Shaanxi2011 |
| KY417150.1 | Bat-CoV_Rs4874 |
| KF569996.1 | Bat-CoV_LYRa11 |
| KY417142.1 | Bat-CoV_As6526 |
| DQ071615.1 | Bat-CoV_Rp3 |
| MK211374.1 | Bat-CoV BC2018 |
| MG772933.1 | Bat-CoV_CVZC45 |
| KT444582.1 | Bat-CoV_WIV16 |
| JX993988.1 | Bat-CoV_Yunnan2011 |
| DQ084200.1 | Bat-CoV_HKU3-3 |
| DQ022305.2 | Bat-CoV_HKU3-1 |
| GQ153544.1 | Bat-CoV_HKU3-9 |
| KJ473813.1 | Bat-CoV BX2013 |
| DQ648857.1 | Bat-CoV BtCoV/279/2005 |
| DQ412043.1 | Bat-CoV_Rm1 |
| KF294457.1 | Bat-CoV_Longquan_140 |
| MG772934.1 | Bat-CoV_CVZXC21 |
| KY352407.1 | Bat-CoV BtKY72 |
| AY394981.1 | BARS_HGZ8L1-A |

| Annotation | lung01 | lung02 | lung03 | lung04 |
|---|---|---|---|---|
| **NC_001552 Murine respirovirus** | **203** | **2002** | **26** | **4769** |
| X00255 Feline sarcoma virus | 40 | 133 | 101 | 72 |
| NC_031450 Parus major densovirus | 2 | 1 | 0 | 0 |
| NC_042001 Arthrobacter phage Molivia | 4 | 2 | 7 | 0 |
| NC_042057 Enterobacteria phage DE3 | 1 | 3.909 | 8.837 | 0 |
| NC_001628 Pseudomonas phage PP7 | 0 | 0 | 0 | 0 |
| NC_038923 Feline sarcoma virus (STRAIN HARDY-Z | 4 | 0 | 1 | 0 |
| NC_001499 Abelson murine leukemia virus | 4 | 1.211 | 0 | 5 |
| NC_029853 Mus musculus mobilized endogenous | 0 | 0 | 4.86 | 0 |
| NC_024909 Caribou feces-associated gemycircular | 0 | 0 | 0 | 0 |
| AF221065 DG-75 Murine leukemia virus | 0 | 0 | 6 | 0 |
| NC_001500 Spleen focus-forming virus | 0 | 0 | 0 | 3 |
| **JX993988 Bat coronavirus Cp/Yunnan2011** | **0** | **0** | **0** | **0** |
| NC_038858 Finkel-Biskis-Jinkins murine sarcoma v | 0.5 | 1 | 4.5 | 0.5 |
| NC_001506 Murine osteosarcoma virus | 0.5 | 1 | 4.5 | 0.5 |
| **KC881005 Bat SARS-like coronavirus RsSHC014** | **0** | **0** | **0** | **0** |
| NC_001501 Moloney murine leukemia virus | 0 | 4.789 | 1 | 4 |
| **KC881006 Bat SARS-like coronavirus Rs3367** | **0** | **0** | **0** | **0** |
| NC_008094 Y73 sarcoma virus | 4 | 0 | 0 | 0 |
| **DQ412043 Bat SARS CoV Rm1/2004** | **0** | **0** | **0** | **0** |
| **GQ153547 Bat SARS coronavirus HKU3-12** | **0** | **0** | **0** | **0** |
| NC_038668 Harvey murine sarcoma virus | 1 | 0 | 0 | 0 |
| NC_001450 Equine infectious anemia virus | 0 | 0 | 0 | 1 |
| NC_001866 Avian myelocytomatosis virus | 0 | 0.5 | 0 | 0 |
| AF033809 Avian myelocytomatosis virus 29 | 0 | 0.5 | 0 | 0 |
| NC_001416 Escherichia virus Lambda | 0 | 0.031 | 1.925 | 0 |
| NC_019488 Salmonella phage RE-2010 | 0 | 0.108 | 0 | 0 |
| X13744 Avian retrovirus IC10 | 0 | 0 | 0 | 2 |
| NC_010463 Salmonella virus Fels2 | 0 | 0 | 0 | 0 |
| NC_024694 Human circovirus VS6600022 | 0 | 0 | 0 | 0 |
| NC_035889 Zika virus | 0 | 0 | 0 | 0 |
| NC_028655 Yersinia phage vB_YenP_AP10 | 0 | 0 | 0 | 0 |
| NC_039044 Feline bocaparvovirus 3 | 0 | 0 | 0 | 0 |
| NC_040671 Adeno-associated virus | 0 | 0 | 0 | 0 |
| NC_028795 Enterobacter phage E-3 | 0 | 0 | 0 | 0 |
| NC_028880 Citrobacter phage phiCFP-1 | 0 | 0 | 0 | 0 |

| lung07 | lung08 | lung09 | lung11 | lung12 | lung13 | lung19 | lymph01 | lymphA01 |
|---|---|---|---|---|---|---|---|---|
| 388 | 1312 | 4375 | 23 | 2 | 0 | 44750 | 1 | 0 |
| 107 | 56 | 3 | 1324 | 7 | 0 | 38 | 21 | 77 |
| 30 | 9 | 0 | 16 | 0 | 14 | 685 | 585 | 1 |
| 83 | 2 | 1 | 24 | 0 | 15 | 1 | 1 | 3 |
| 0.231 | 0 | 0 | 5 | 0 | 269.987 | 8.638 | 1 | 30.906 |
| 0 | 0 | 0 | 0 | 0 | 0 | 131 | 0 | 0 |
| 0 | 0 | 64 | 17 | 0 | 0 | 0 | 0 | 0 |
| 2 | 0 | 57 | 1 | 0 | 0 | 0 | 0 | 0 |
| 5 | 0 | 0 | 0 | 0 | 0 | 8 | 8 | 4 |
| 1 | 0 | 0 | 0 | 0 | 0 | 0 | 64 | 0 |
| 7.804 | 0 | 0 | 0 | 2 | 0 | 3 | 2.639 | 2 |
| 0 | 0 | 0 | 1 | 0 | 0 | 1 | 1.361 | 0 |
| **6.745** | **43.936** | **0.039** | **0** | **0** | **0** | **0** | **0** | **0** |
| 3.5 | 0 | 0 | 1.5 | 0 | 0 | 7.5 | 2 | 2.5 |
| 3.5 | 0 | 0 | 1.5 | 0 | 0 | 7.5 | 2 | 2.5 |
| **5.211** | **31.405** | **0** | **0.075** | **0** | **0** | **0** | **0** | **0** |
| 0 | 4 | 0 | 0 | 0 | 0 | 5 | 0 | 0 |
| **0.036** | **18.32** | **1.938** | **0.075** | **0** | **0** | **0** | **0** | **0** |
| 0 | 0 | 10 | 0 | 0 | 0 | 0 | 0 | 0 |
| **2.03** | **11.969** | **0.144** | **0** | **0** | **0** | **0** | **0** | **0** |
| **6.839** | **4.162** | **0.143** | **0** | **0** | **0** | **0** | **0** | **0** |
| 0 | 0 | 6 | 1 | 0 | 0 | 0 | 0 | 0 |
| 0 | 0 | 0 | 0 | 0 | 0 | 0 | 0 | 2 |
| 0 | 0 | 3.5 | 1 | 0 | 0 | 0 | 1 | 0 |
| 0 | 0 | 3.5 | 1 | 0 | 0 | 0 | 1 | 0 |
| 0.26 | 0 | 0 | 0 | 0 | 0.004 | 0.096 | 0 | 0.04 |
| 0 | 0 | 1 | 0 | 0 | 0 | 1.98 | 0 | 0 |
| 0 | 1 | 1 | 0 | 0 | 0 | 0 | 0 | 0 |
| 0 | 0 | 0 | 0 | 0 | 0 | 0 | 0 | 0 |
| 0 | 0 | 0 | 0 | 0 | 0 | 0 | 0 | 0 |
| 0 | 0 | 0 | 0 | 0 | 0 | 0 | 0 | 0 |
| 0 | 0 | 0 | 0 | 0 | 0 | 0 | 0 | 0 |
| 0 | 0 | 0 | 0 | 0 | 0 | 0 | 0 | 0 |
| 0 | 0 | 0 | 0 | 0 | 0 | 0 | 0 | 0 |
| 0 | 0 | 0 | 0 | 0 | 0 | 0 | 0 | 0 |
| 0 | 0 | 0 | 0 | 0 | 0 | 0 | 0 | 0 |

| spleen01 | spleen02 | spleen03 | spleen04 | spleen08 | spleen11 | spleen12 | spleen19 | Lineage |
|---|---|---|---|---|---|---|---|---|
| 0 | 11 | 0 | 15 | 12 | 20 | 0 | 0 | Viruses; Ri |
| 0 | 7 | 2 | 3 | 7 | 5 | 3 | 5 | Viruses; Or |
| 0 | 0 | 26 | 8 | 0 | 2 | 0 | 72 | Viruses; Pa |
| 0 | 0 | 0 | 0 | 12 | 8 | 6 | 423 | Viruses; Ca |
| 1 | 10 | 32 | 6.596 | 4 | 6 | 20 | 6.889 | Viruses; Ca |
| 0 | 0 | 0 | 0 | 0 | 0 | 0 | 0 | Viruses; Rib |
| 0 | 0 | 0 | 1 | 0 | 0 | 1 | 0 | Viruses; Or |
| 0 | 1 | 0 | 0 | 1 | 0 | 0 | 0 | Viruses; Or |
| 0 | 9 | 0 | 13.833 | 3 | 12 | 0 | 0 | Viruses; Or |
| 0 | 0 | 0 | 0 | 0 | 0 | 0 | 0 | Viruses; Ge |
| 3.655 | 8.348 | 0 | 4.851 | 11 | 5.215 | 1 | 0 | Viruses; Or |
| 6.345 | 1.247 | 1 | 0 | 0 | 2.785 | 0 | 35 | Viruses; Or |
| 0 | 0 | 0 | 0 | 0 | 0 | 0 | 0 | Viruses; Ri |
| 0 | 2 | 0 | 9.5 | 6.5 | 7 | 0 | 0 | Viruses; Or |
| 0 | 2 | 0 | 9.5 | 6.5 | 7 | 0 | 0 | Viruses; Or |
| 0 | 0 | 0 | 0 | 0 | 0 | 0 | 0 | Viruses; Ri |
| 0 | 1 | 0 | 2 | 0 | 0 | 0 | 0 | Viruses; Or |
| 0 | 0 | 0 | 0 | 0 | 0 | 0 | 0 | Viruses; Ri |
| 0 | 1 | 0 | 0 | 0 | 0 | 0 | 0 | Viruses; Or |
| 0 | 0 | 0 | 0 | 0 | 0 | 0 | 0 | Viruses; Ri |
| 0 | 0 | 0 | 0 | 0 | 0 | 0 | 0 | Viruses; Ri |
| 0 | 0 | 0 | 0 | 0 | 1 | 0 | 0 | Viruses; Or |
| 0 | 0 | 3 | 0 | 1 | 1 | 0 | 0 | Viruses; Or |
| 0 | 0 | 0 | 0 | 0 | 0 | 0 | 0 | Viruses; Or |
| 0 | 0 | 0 | 0 | 0 | 0 | 0 | 0 | Viruses; Or |
| 0 | 0.822 | 0 | 1.046 | 0 | 0 | 0 | 0.038 | Viruses; Ca |
| 0 | 0 | 0.16 | 0 | 0 | 0 | 0 | 1 | Viruses; Ca |
| 0 | 0 | 0 | 0 | 0 | 0 | 0 | 0 | Viruses; Or |
| 0 | 0 | 0.84 | 0 | 0 | 0 | 0 | 0 | Viruses; Ca |
| 0 | 0 | 0 | 0 | 0 | 0 | 0 | 0 | Viruses; Cir |
| 0 | 0 | 0 | 0 | 0 | 0 | 0 | 0 | Viruses; Rib |
| 0 | 0 | 0 | 0 | 0 | 0 | 0 | 0 | Viruses; Ca |
| 0 | 0 | 0 | 0 | 0 | 0 | 0 | 0 | Viruses; Pa |
| 0 | 0 | 0 | 0 | 0 | 0 | 0 | 0 | Viruses; Pa |
| 0 | 0 | 0 | 0 | 0 | 0 | 0 | 0 | Viruses; Ca |
| 0 | 0 | 0 | 0 | 0 | 0 | 0 | 0 | Viruses; Ca |

**iboviria; Negarnaviricota; Haploviricotina; Monjiviricetes; Mononegavirales; Paramyxoviridae;**
tervirales; Retroviridae; Orthoretrovirinae; Gammaretrovirus; Feline leukemia virus
rvoviridae; Densovirinae; unclassified Densovirinae
udovirales; Siphoviridae; Amigovirus; Arthrobacteria virus Molivia
udovirales; Siphoviridae; Lambdavirus; Escherichia virus DE3
poviria; Leviviridae; unclassified Leviviridae
tervirales; Retroviridae; Orthoretrovirinae; Gammaretrovirus; Hardy-Zuckerman feline sarcoma virus
tervirales; Retroviridae; Orthoretrovirinae; Gammaretrovirus; unclassified Gammaretrovirus
tervirales; Retroviridae; Orthoretrovirinae; Gammaretrovirus; unclassified Gammaretrovirus
Organisma 9870
tervirales; Retroviridae; Orthoretrovirinae; Gammaretrovirus; Murine leukemia virus; unclassified Mu
tervirales; Retroviridae; Orthoretrovirinae; Gammaretrovirus; unclassified Gammaretrovirus
**iboviria; Nidovirales; Cornidovirineae; Coronaviridae; Orthocoronavirinae; Betacoronavirus; Sa**
tervirales; Retroviridae; Orthoretrovirinae; Gammaretrovirus
tervirales; Retroviridae; Orthoretrovirinae; Gammaretrovirus; unclassified Gammaretrovirus
**iboviria; Nidovirales; Cornidovirineae; Coronaviridae; Orthocoronavirinae; Betacoronavirus; Sa**
tervirales; Retroviridae; Orthoretrovirinae; Gammaretrovirus; Murine leukemia virus
**iboviria; Nidovirales; Cornidovirineae; Coronaviridae; Orthocoronavirinae; Betacoronavirus; Sa**
tervirales; Retroviridae; Orthoretrovirinae; Alpharetrovirus
**iboviria; Nidovirales; Cornidovirineae; Coronaviridae; Orthocoronavirinae; Betacoronavirus; Sa**
**iboviria; Nidovirales; Cornidovirineae; Coronaviridae; Orthocoronavirinae; Betacoronavirus; Sa**
tervirales; Retroviridae; Orthoretrovirinae; Gammaretrovirus
tervirales; Retroviridae; Orthoretrovirinae; Lentivirus
tervirales; Retroviridae; Orthoretrovirinae; Alpharetrovirus; unclassified Alpharetrovirus
tervirales; Retroviridae; Orthoretrovirinae; Alpharetrovirus
udovirales; Siphoviridae; Lambdavirus
udovirales; Myoviridae; Peduovirinae; Peduovirus; unclassified Peduovirus
tervirales; Retroviridae; Orthoretrovirinae; Alpharetrovirus; unclassified Alpharetrovirus
udovirales; Myoviridae; Peduovirinae; Peduovirus
coviridae; Circovirus; unclassified Circovirus
poviria; Flaviviridae; Flavivirus
udovirales; Podoviridae; Autographivirinae; Teseptimavirus; unclassified Teseptimavirus
rvoviridae; Parvovirinae; Bocaparvovirus; Carnivore bocaparvovirus 5
rvoviridae; Parvovirinae; Dependoparvovirus; unclassified Dependoparvovirus
udovirales; Podoviridae; Autographivirinae; Teseptimavirus; unclassified Teseptimavirus
udovirales; Podoviridae; Autographivirinae; Teseptimavirus; unclassified Teseptimavirus

Orthoparamyxovirinae; Respirovirus

;

ırine leukemia virus

rbecovirus; Severe acute respiratory syndrome-related coronavirus

rbecovirus; Severe acute respiratory syndrome-related coronavirus

rbecovirus; Severe acute respiratory syndrome-related coronavirus

rbecovirus; Severe acute respiratory syndrome-related coronavirus
rbecovirus; Severe acute respiratory syndrome-related coronavirus; Bat SARS coronavirus HK

U3

### BLASTN 2.9.0+
### Query: 2019-nCoV NC_045512.2 Wuhan seafood market pneumonia virus isolate Wuhan-Hu-1,
### Database: User specified sequence set (Input: V2-M_Cov.contigs-convert.fa)
### Fields: query acc.ver, subject acc.ver, % identity, alignment length, mismatches, gap opens, q. star
### 53 hits found

| | | | | | | | |
|---|---|---|---|---|---|---|---|
| 2019-nCo\789-1-k14 | 91.095 | 3369 | 298 | 2 | 14707 | 18074 | 1 |
| 2019-nCo\789-1-k14 | 92.923 | 2812 | 199 | 0 | 11849 | 14660 | 2872 |
| 2019-nCo\789-1-k14 | 94.231 | 52 | 3 | 0 | 14063 | 14114 | 52 |
| 2019-nCo\789-1-k14 | 94.687 | 2202 | 117 | 0 | 27254 | 29455 | 1 |
| 2019-nCo\789-1-k14 | 98.739 | 238 | 3 | 0 | 29485 | 29722 | 2430 |
| 2019-nCo\789-1-k14 | 87.28 | 2107 | 231 | 26 | 19890 | 21978 | 2278 |
| 2019-nCo\789-1-k14 | 91.522 | 1156 | 94 | 4 | 233 | 1386 | 4 |
| 2019-nCo\789-1-k14 | 88 | 1325 | 157 | 2 | 18277 | 19600 | 1324 |
| 2019-nCo\789-1-k14 | 90.617 | 1151 | 108 | 0 | 23926 | 25076 | 1 |
| 2019-nCo\789-1-k14 | 91.284 | 1044 | 91 | 0 | 9176 | 10219 | 1044 |
| 2019-nCo\789-1-k14 | 94.779 | 747 | 39 | 0 | 26171 | 26917 | 1067 |
| 2019-nCo\789-1-k14 | 96.622 | 296 | 10 | 0 | 26016 | 26311 | 296 |
| 2019-nCo\789-1-k14 | 100 | 37 | 0 | 0 | 26434 | 26470 | 289 |
| 2019-nCo\789-1-k14 | 89.055 | 804 | 88 | 0 | 5050 | 5853 | 873 |
| 2019-nCo\M-1-k141_ | 91.006 | 656 | 57 | 2 | 674 | 1328 | 3 |
| 2019-nCo\789-1-k14 | 94.251 | 574 | 33 | 0 | 11289 | 11862 | 574 |
| 2019-nCo\789-1-k14 | 85.644 | 815 | 117 | 0 | 22382 | 23196 | 4 |
| 2019-nCo\k141_1140 | 93.974 | 531 | 32 | 0 | 11895 | 12425 | 1 |
| 2019-nCo\789-3-k14 | 95.397 | 478 | 22 | 0 | 28704 | 29181 | 720 |
| 2019-nCo\789-3-k14 | 95.146 | 103 | 5 | 0 | 28627 | 28729 | 237 |
| 2019-nCo\789-3-k14 | 91.089 | 101 | 9 | 0 | 27590 | 27690 | 101 |
| 2019-nCo\k141_1571 | 93.387 | 499 | 33 | 0 | 17074 | 17572 | 546 |
| 2019-nCo\789-3-k14 | 93.387 | 499 | 33 | 0 | 17074 | 17572 | 546 |
| 2019-nCo\789-2-k14 | 93.387 | 499 | 33 | 0 | 17074 | 17572 | 546 |
| 2019-nCo\M-2-k141_ | 90.963 | 509 | 44 | 2 | 626 | 1133 | 171 |
| 2019-nCo\k141_8112 | 95 | 420 | 21 | 0 | 28949 | 29368 | 1 |
| 2019-nCo\k141_9061 | 91.277 | 470 | 41 | 0 | 7295 | 7764 | 470 |
| 2019-nCo\k141_1217 | 94.293 | 403 | 23 | 0 | 26849 | 27251 | 403 |
| 2019-nCo\k141_8467 | 98.551 | 345 | 5 | 0 | 29483 | 29827 | 1 |
| 2019-nCo\789-2-k14 | 96.552 | 348 | 12 | 0 | 28627 | 28974 | 569 |
| 2019-nCo\789-2-k14 | 94.098 | 305 | 18 | 0 | 28877 | 29181 | 259 |
| 2019-nCo\789-2-k14 | 98.718 | 156 | 2 | 0 | 29213 | 29368 | 86 |
| 2019-nCo\789-2-k14 | 91.549 | 71 | 6 | 0 | 29294 | 29364 | 1 |
| 2019-nCo\k141_1241 | 89.286 | 448 | 48 | 0 | 4293 | 4740 | 1 |
| 2019-nCo\789-2-k14 | 90.704 | 398 | 36 | 1 | 25728 | 26125 | 84 |
| 2019-nCo\789-2-k14 | 89.057 | 265 | 29 | 0 | 26750 | 27014 | 738 |
| 2019-nCo\789-2-k14 | 92.683 | 82 | 6 | 0 | 25848 | 25929 | 1 |
| 2019-nCo\k141_8691 | 91.214 | 387 | 34 | 0 | 8437 | 8823 | 1 |
| 2019-nCo\k141_4684 | 96.774 | 310 | 10 | 0 | 28338 | 28647 | 8 |
| 2019-nCo\k141_1020 | 91.549 | 355 | 30 | 0 | 12572 | 12926 | 446 |
| 2019-nCo\k141_7268 | 88.832 | 394 | 44 | 0 | 1571 | 1964 | 2 |
| 2019-nCo\789-2-k14 | 86.247 | 429 | 59 | 0 | 18957 | 19385 | 1 |
| 2019-nCo\k141_1403 | 87.812 | 361 | 32 | 1 | 23550 | 23910 | 1 |
| 2019-nCo\k141_8774 | 86.216 | 370 | 51 | 0 | 18957 | 19326 | 370 |
| 2019-nCo\k141_7634 | 87.106 | 349 | 44 | 1 | 16415 | 16762 | 525 |
| 2019-nCo\k141_7634 | 95.726 | 117 | 5 | 0 | 16251 | 16367 | 162 |
| 2019-nCo\789-3-k14 | 87.106 | 349 | 44 | 1 | 16415 | 16762 | 525 |
| 2019-nCo\789-3-k14 | 95.726 | 117 | 5 | 0 | 16251 | 16367 | 162 |
| 2019-nCo\789-2-k14 | 87.106 | 349 | 44 | 1 | 16415 | 16762 | 525 |
| 2019-nCo\789-2-k14 | 95.726 | 117 | 5 | 0 | 16251 | 16367 | 162 |
| 2019-nCo\k141_1282 | 95.023 | 221 | 11 | 0 | 27299 | 27519 | 1 |
| 2019-nCo\k141_1525 | 91.388 | 209 | 18 | 0 | 28009 | 28217 | 100 |
| 2019-nCo\k141_1525 | 98.684 | 76 | 1 | 0 | 28241 | 28316 | 21 |

complete genome

t, q. end, s. start, s. end, evalue, bit score

| | | |
|---|---|---|
| 3368 | 0 | 4558 |
| 61 | 0 | 4091 |
| 1 | 5.45E-16 | 80.5 |
| 2202 | 0 | 3419 |
| 2193 | 2.18E-119 | 424 |
| 191 | 0 | 2372 |
| 1157 | 0 | 1589 |
| 1 | 0 | 1565 |
| 1151 | 0 | 1528 |
| 1 | 0 | 1424 |
| 321 | 0 | 1164 |
| 1 | 5.90E-140 | 492 |
| 325 | 1.18E-12 | 69.4 |
| 70 | 0 | 998 |
| 657 | 0 | 883 |
| 1 | 0 | 878 |
| 818 | 0 | 857 |
| 531 | 0 | 804 |
| 243 | 0 | 761 |
| 135 | 5.26E-41 | 163 |
| 1 | 3.19E-33 | 137 |
| 48 | 0 | 739 |
| 48 | 0 | 739 |
| 48 | 0 | 739 |
| 678 | 0 | 684 |
| 420 | 0 | 660 |
| 1 | 0 | 641 |
| 1 | 9.33E-178 | 617 |
| 345 | 1.56E-175 | 610 |
| 916 | 1.58E-165 | 577 |
| 563 | 1.29E-131 | 464 |
| 241 | 1.80E-75 | 278 |
| 71 | 1.50E-21 | 99 |
| 448 | 4.44E-161 | 562 |
| 480 | 4.50E-151 | 529 |
| 474 | 4.90E-91 | 329 |
| 82 | 1.15E-27 | 119 |
| 387 | 1.62E-150 | 527 |
| 317 | 9.74E-148 | 518 |
| 92 | 2.12E-139 | 490 |
| 395 | 9.88E-138 | 484 |
| 429 | 3.58E-132 | 466 |
| 349 | 4.73E-116 | 412 |
| 1 | 1.02E-112 | 401 |
| 177 | 1.71E-110 | 394 |
| 46 | 8.67E-49 | 189 |
| 177 | 1.71E-110 | 394 |
| 46 | 8.67E-49 | 189 |
| 177 | 1.71E-110 | 394 |
| 46 | 8.67E-49 | 189 |
| 221 | 1.35E-96 | 348 |
| 308 | 2.99E-78 | 287 |
| 96 | 1.15E-32 | 135 |

### BLASTX 2.9.0+
### Query: Pan_CoV
### Database: User specified sequence set (Input: ../NC_045512.2.txt)
### Fields: query acc.ver, subject acc.ver, % identity, alignment length, mismatches, gap opens, q.
### 34 hits found

| | | | | | | | |
|---|---|---|---|---|---|---|---|
| Pan_CoV | lcl\|NC_045! | 94.722 | 2084 | 71 | 2 | 14173 | 20421 |
| Pan_CoV | lcl\|NC_045! | 72.148 | 1490 | 337 | 5 | 4142 | 8461 |
| Pan_CoV | lcl\|NC_045! | 94.798 | 769 | 3 | 1 | 11873 | 14179 |
| Pan_CoV | lcl\|NC_045! | 91.808 | 647 | 53 | 0 | 8894 | 10834 |
| Pan_CoV | lcl\|NC_045! | 98.407 | 565 | 9 | 0 | 21451 | 23145 |
| Pan_CoV | lcl\|NC_045! | 77.059 | 510 | 95 | 4 | 1436 | 2932 |
| Pan_CoV | lcl\|NC_045! | 95.2 | 375 | 18 | 0 | 146 | 1270 |
| Pan_CoV | lcl\|NC_045! | 88.957 | 163 | 15 | 1 | 20521 | 21000 |
| Pan_CoV | lcl\|NC_045! | 89.441 | 161 | 17 | 0 | 3455 | 3937 |
| Pan_CoV | lcl\|NC_045! | 98.462 | 65 | 1 | 0 | 21007 | 21201 |
| Pan_CoV | lcl\|NC_045! | 100 | 12 | 0 | 0 | 20969 | 21004 |
| Pan_CoV | lcl\|NC_045! | 89.394 | 66 | 7 | 0 | 2108 | 2305 |
| Pan_CoV | lcl\|NC_045! | 100 | 49 | 0 | 0 | 11351 | 11497 |
| Pan_CoV | lcl\|NC_045! | 91.667 | 24 | 2 | 0 | 1334 | 1263 |
| Pan_CoV | lcl\|NC_045! | 72.148 | 1490 | 337 | 5 | 4142 | 8461 |
| Pan_CoV | lcl\|NC_045! | 94.825 | 773 | 3 | 1 | 11873 | 14191 |
| Pan_CoV | lcl\|NC_045! | 91.808 | 647 | 53 | 0 | 8894 | 10834 |
| Pan_CoV | lcl\|NC_045! | 77.059 | 510 | 95 | 4 | 1436 | 2932 |
| Pan_CoV | lcl\|NC_045! | 95.2 | 375 | 18 | 0 | 146 | 1270 |
| Pan_CoV | lcl\|NC_045! | 89.441 | 161 | 17 | 0 | 3455 | 3937 |
| Pan_CoV | lcl\|NC_045! | 89.394 | 66 | 7 | 0 | 2108 | 2305 |
| Pan_CoV | lcl\|NC_045! | 100 | 49 | 0 | 0 | 11351 | 11497 |
| Pan_CoV | lcl\|NC_045! | 91.667 | 24 | 2 | 0 | 1334 | 1263 |
| Pan_CoV | lcl\|NC_045! | 96.673 | 511 | 13 | 1 | 25129 | 26649 |
| Pan_CoV | lcl\|NC_045! | 78.189 | 541 | 113 | 3 | 23209 | 24816 |
| Pan_CoV | lcl\|NC_045! | 97.462 | 394 | 10 | 0 | 29857 | 31038 |
| Pan_CoV | lcl\|NC_045! | 98.198 | 222 | 4 | 0 | 28106 | 28771 |
| Pan_CoV | lcl\|NC_045! | 96.226 | 212 | 8 | 0 | 27165 | 27800 |
| Pan_CoV | lcl\|NC_045! | 94.215 | 121 | 7 | 0 | 29477 | 29839 |
| Pan_CoV | lcl\|NC_045! | 97.521 | 121 | 3 | 0 | 28977 | 29339 |
| Pan_CoV | lcl\|NC_045! | 96.721 | 61 | 2 | 0 | 28785 | 28967 |
| Pan_CoV | lcl\|NC_045! | 100 | 75 | 0 | 0 | 27828 | 28052 |
| Pan_CoV | lcl\|NC_045! | 97.368 | 38 | 1 | 0 | 31193 | 31080 |
| Pan_CoV | lcl\|NC_045! | 97.368 | 38 | 1 | 0 | 31441 | 31554 |

start, q. end, s. start, s. end, evalue, bit score

| | | | |
|---|---|---|---|
| 4400 | 6445 | 0 | 4148 |
| 1044 | 2505 | 0 | 1943 |
| 3670 | 4401 | 0 | 1400 |
| 2677 | 3323 | 0 | 1232 |
| 6527 | 7091 | 0 | 1159 |
| 137 | 635 | 0 | 792 |
| 1 | 375 | 0 | 784 |
| 6232 | 6394 | 8.74E-88 | 315 |
| 810 | 970 | 1.24E-71 | 262 |
| 6164 | 6228 | 6.98E-42 | 149 |
| 6151 | 6162 | 6.98E-42 | 35.8 |
| 117 | 182 | 7.65E-33 | 133 |
| 3496 | 3544 | 3.91E-24 | 104 |
| 129 | 152 | 3.32E-08 | 52 |
| 1044 | 2505 | 0 | 1951 |
| 3670 | 4405 | 0 | 1414 |
| 2677 | 3323 | 0 | 1237 |
| 137 | 635 | 0 | 795 |
| 1 | 375 | 0 | 787 |
| 810 | 970 | 6.35E-72 | 263 |
| 117 | 182 | 4.79E-33 | 134 |
| 3496 | 3544 | 3.01E-24 | 105 |
| 129 | 152 | 3.26E-08 | 52 |
| 658 | 1168 | 0 | 1006 |
| 13 | 553 | 0 | 888 |
| 1 | 394 | 0 | 590 |
| 1 | 222 | 1.85E-145 | 444 |
| 64 | 275 | 1.84E-127 | 395 |
| 1 | 121 | 6.48E-76 | 241 |
| 1 | 121 | 1.36E-68 | 219 |
| 1 | 61 | 6.40E-34 | 118 |
| 1 | 75 | 4.69E-27 | 99.8 |
| 1 | 38 | 1.49E-19 | 77 |
| 1 | 38 | 1.49E-19 | 77 |
